# Identification of qPCR reference genes suitable for normalising gene expression in the developing mouse embryo

**DOI:** 10.1101/2020.10.29.360461

**Authors:** John C.W. Hildyard, Dominic J. Wells, Richard J. Piercy

## Abstract

Mammalian embryogenesis is an intricate, tightly orchestrated process. Progression from zygote through somitogenesis and on to organogenesis and maturity involves many interacting cell types and multiple differentiating cell lineages. Quantitative PCR analysis of gene expression in the developing embryo is a valuable tool for deciphering these interactions and tracing lineages, but normalisation of qPCR data to stably expressed reference genes is essential. Patterns of gene expression change globally and dramatically as embryonic development proceeds, rendering identification of appropriate reference genes challenging at both the whole embryo- and individual tissue-level. We have investigated expression stability in mouse embryos from mid to late gestation (E11.5–E18.5), both at the whole-embryo level, and within more restricted tissue domains (head, developing forelimb), using 15 candidate reference genes (*ACTB, 18S, SDHA, GAPDH, HTATSF1, CDC40, RPL13A, CSNK2A2, AP3D1, HPRT1, CYC1, EIF4A, UBC, B2M* and *PAK1IP1*), and four complementary algorithms (geNorm, Normfinder, Bestkeeper and deltaCt). Unexpectedly, all methods suggest that many genes within our candidate panel are acceptable references, and despite disagreement over highest-scoring candidates, *AP3D1*, *RPL14A* and *PAK1IP1* are the strongest performing genes overall. Conversely, *HPRT1* and *B2M* are consistently poor choices: these genes show strong developmental regulation. We further show that use of *AP3D1*, *RPL13A* and *PAK1IP1* can reveal subtle patterns of developmental expression even in genes ostensibly ranked as acceptable (*CDC40*, *HTATSF1*), and thus these three represent universally suitable reference genes for the mouse embryo.

## Introduction

The mouse (*Mus musculus*) is the principal model organism for the study of mammalian embryonic development, offering large litter sizes and a predictable gestation period (mating in this species can moreover be timed with relative ease). The stages of mouse embryonic development are thus well documented [1, 2], and given the conserved nature of mammalian embryogenesis, can also be mapped across mammalian species despite gestation times differing by more than an order of magnitude (20 days in the mouse, 270 days in the human, up to 645 days in the African elephant *Loxodonta*) [3].

The earliest stages of development establish fundamental morphological patterns: blastocyst formation, implantation, establishment of polarity occur with the first five days (Embryonic days 1-5, or E1-E5) [4], with gastrulation and development of the primitive streak following (E6-E8.5) [5]. Intermediate stages (E8.5-E10.5) feature the cyclical ‘clock and wave’ of somitogenesis [6], the turning of the embryo, neural tube closure and the laying down of organ precursor cell lineages, while the later stages (E11.5 onward) involve maturation and development of those organs [1, 2]. During this later period (from E11.5 to E18.5, Fig 1) almost all limb development occurs, from a primitive limb bud to an essentially mature state complete with ossifying bones and fully-defined joints and digits [7–9], and similarly almost all skeletal muscle is laid down (both primary and secondary myogenesis) [10, 11]. The head undergoes a broad panel of changes, with the brain enlarging and maturing [12], the palate closing and teeth forming [13], the ears emerging, and during this period the eye progresses from a simple indentation to a fully encapsulated globe (which is subsequently sheltered beneath fused eyelids) [14]. More globally, this later period also spans substantial changes in haematopoiesis (with the source of blood cells switching from the yolk sac to the liver, and then subsequently to the bone marrow) [15], maturation of the chambers of the heart [16], gradual replacement of the mesonephros with the metanephros [17], and establishment of the skin barrier [1, 18].

**Fig 1:**
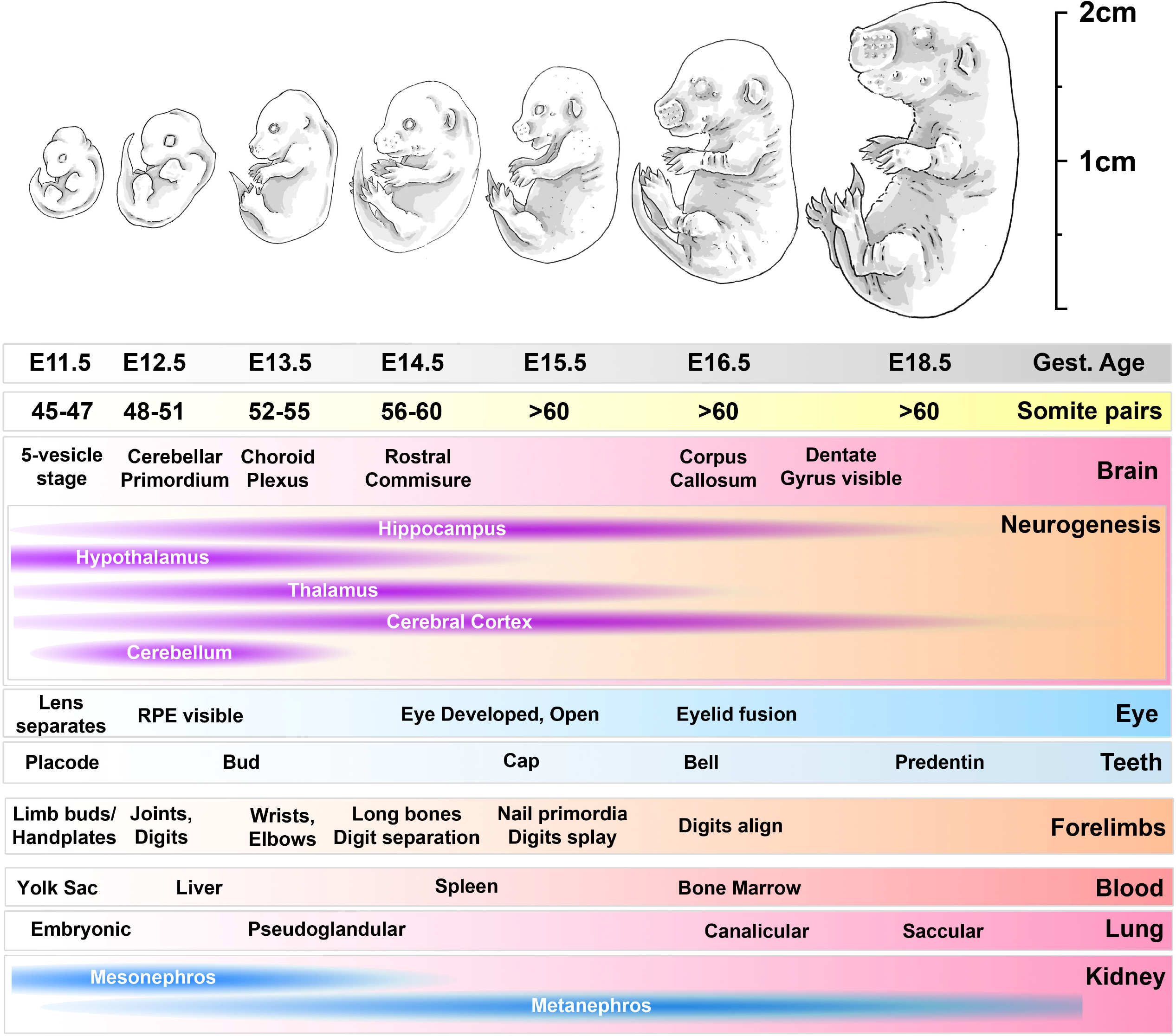
developmental timeline of mouse embryos used in this study Progress from E11.5 to E18.5 involves marked increases in size and mass and progressive accumulation of somite pairs. Within the head, multiple defined brain structures emerge over this period, and neurogenesis peaks in different brain regions (as indicated). The eye matures, as do teeth. Within the forelimbs, development proceeds from primitive limb bud through to joint and digit formation, maturation and ossification of long bones and emergence of nail primordia. Within the body, the source of haematopoiesis shifts from the yolk sac to the liver, and thence transiently to the spleen before establishing in the bone marrow; the lungs mature through primitive pseudoglandular stages to saccular morphology; the mesonephros of the kidney degenerates and is replaced with the final metanephros. RPE: retinal pigmented epithelium

Although the morphological and developmental changes during embryogenesis are well-characterised at the histological level, understanding at the molecular level is less comprehensive. The interactions of conflicting growth factors, signalling cascades and downstream transcriptional programs play a critical role in coordination of development, but the diversity of cell types and the often transient nature of such interactions renders these processes challenging to investigate. More focussed data can be obtained by careful isolation of specific embryonic tissues, however not all tissues are tractable to such an approach, and restricting analysis in this fashion loses wider developmental context. Recent technological advances have permitted elegant studies of embryonic gene expression down to single-cell level [19, 20], suggesting that detailed study of the interactions between individual cell lineages might now be possible, however such approaches also represent a substantial undertaking both in time and resources. Furthermore, these approaches are not without limitation: the whole transcriptome nature of the method typically favours breadth over depth [21, 22]: large-scale changes involving many gene components will be mapped well, while more specific, smaller-scale processes can be missed. In particular, constraints in read depth mean low abundance transcripts can be under-represented in such datasets [23], and thus more conventional methods remain vital research tools.

Measurement of gene expression via qPCR (using cDNA prepared from extracted RNA) is a comparatively inexpensive and flexible means of measuring expression of specific gene targets, including rare transcripts expressed at low levels or restricted to minority cell populations. Given the quantitative nature of this technique, normalisation of expression data is essential: RNA extraction efficiency, RNA integrity and cDNA synthesis efficiency all represent sources of variability that can potentially obscure genuine changes in gene expression, or create the appearance of change where none exists (best summarised in the MIQE guidelines, [24]). Normalisation can be conducted using reference genes: genes known to be stably expressed under the conditions studied. RNA is an inherently labile molecule, however, and cellular turnover at the transcript level is often both more dramatic, and more rapid, than at the protein level. Genes considered broadly appropriate for protein normalisation such as *GAPDH* and *ACTB* (beta-actin) are historically popular choices, but we and others have shown these often perform poorly as references [25–31]. 18S ribosomal RNA is also used widely, but this necessitates use of random priming in cDNA preparation (rRNA lacks polyA tails, thus oligo dT priming will not reverse transcribe *18S*). Synthesis and degradation of ribosomal RNA also differs from that of mRNA [32], and the sheer abundance of rRNA sequences could mask marked changes in the mRNA pool (ribosomes represent ~80-90% of total cellular RNA, while mRNA accounts for ~5%, thus mRNA levels could change by a factor of two or more without significantly altering measured rRNA content). Crucially, reference genes suitable for one comparative scenario are not guaranteed to be appropriate for another, and identification of reference genes appropriate for the conditions studied represents a key step in any qPCR-based investigation. Identification of appropriate reference genes *a priori* is challenging, and various mathematical approaches exist: the geNorm [33], Normfinder [34], Bestkeeper [35] and deltaCt [36] methods all require a representative collection of cDNA samples, and a broad panel of candidate reference genes, but each assesses suitability via subtly different criteria (see supplementary data for an overview). Combining these complementary approaches increases the power of investigations: individual rankings might differ between methods, but truly strong candidates should consistently score highly regardless of assessment method (and discrepancies between methods can moreover highlight interesting biological information, as reported previously [26, 37]).

The developing embryo represents an especially dynamic transcriptional environment, transcriptional changes are likely to be dramatic as development proceeds, making determination of reference genes especially critical; such developmental changes might be even more profound in a mammal with a short gestation like the mouse. Several studies have investigated appropriate references for earlier developmental stages, with a focus on early morula/blastocyst formation [38–40], or progression from blastocyst (E3.5) to mid organogenesis (E11.5) [41, 42], though these studies have produced conflicting results, with *ACTB* scoring highly in some, but poorly in others (some investigators have reported that embryonic reference gene suitability might even vary by mouse strain [39]). Less attention has been dedicated to later stages of embryogenesis, where cell and organ lineages are more established, and developmental changes become consequently more focussed and tissue-specific. Several studies have been conducted on specific organs (such as the developing gonads [43], heart [44] or thymus [45]), but no specific study has addressed reference genes appropriate for normalisation at the whole embryo level at these later stages.

We have thus investigated reference genes appropriate for normalising gene expression in whole mouse embryos collected at E11.5, E12.5, E13.5, E14.5, E15.5, E16.5 and E18.5, spanning development from mid to late gestation. Given the marked developmental changes that occur within the head and the forelimbs over this period (Fig 1), we have also determined reference genes appropriate for these specific tissues (at E13.5, E16.5 and E18.5). Our collection of samples is substantial (N=3-5 per time point) and we have used a broad panel of 15 candidate reference genes (*ACTB, 18S, SDHA, GAPDH, HTATSF1, CDC40, RPL13A, CSNK2A2, AP3D1, HPRT1, CYC1, EIF4A, UBC, B2M* and *PAK1IP1* - Table 1). This panel includes candidates widely used historically (*18S*, *GAPDH*, *ACTB*), those that we have shown be strong references in mouse skeletal muscle (*CSNK2A2*, *AP3D1*, *RPL13A*) [25, 37], and also shown by others to be viable references in rodent brains (*UBC*, *HPRT1*, *RPL13A, 18S*) [46, 47]. To maximise the power of our study, we have assessed reference gene suitability using all four algorithms described above (geNorm, deltaCt, BestKeeper and Normfinder).

**Table 1:** list of candidate genes and their full names/functions

| Gene name | Full name/function |
| --- | --- |
| ACTB | Beta-Actin: cytoskeletal component |
| 18S | 18S ribosomal RNA: small ribosomal subunit |
| SDHA | Succinate Dehydrogenase subunit A: mitochondrial complex II |
| GAPDH | Glyceraldehyde 3-Phosphate Dehydrogenase: glycolysis pathway |
| HTATSF1 | HIV-1 Tat-specific factor 1: transcriptional elongation/splicing factor |
| CDC40 | Cell Division Cycle 40: cell cycle/spliceosome factor |
| RPL13a | Ribosomal Protein L13: component of large ribosomal subunit |
| CSNK2A2 | Casein Kinase 2 Alpha 2: subunit of casein kinase |
| AP3D1 | Adaptor Related Protein Complex 3 Subunit Delta 1: vesicle trafficking |
| HPRT1 | Hypoxanthine Phosphoribosyltransferase 1: purine salvage pathway |
| CYC1 | Cytochrome C1: mitochondrial complex III |
| EIF4A | Eukaryotic Initiation Factor 4A: helicase component of translation initiation complex |
| UBC | Ubiquitin C |
| B2M | Beta 2 Microglobulin: component of MHC class 1 complex |
| PAK1IP1 | PAK1 Interacting Protein: negative regulator of PAK1 kinase |

## Methods

### Ethics statement

A total of 70 mouse embryos (strain C57BL/10) were obtained post-mortem from 10 pregnant females: 35 embryos were used for this study (additional embryos were collected for a separate study). All mice were bred under UK Home Office Project Licence, approved by the Royal Veterinary College Animal Welfare and Ethical Review Body. Mice were held in individually ventilated cages in a minimal disease unit at an average 21°C in a 12 hours light/ 12 hours dark light cycle with food and water provided ad-lib. Mating trios (one male, 2 females) were set up and monitored each morning until all females had been mated (as shown by a vaginal plug). All pregnant females were killed by cervical dislocation at the required gestational age and all embryos were killed via hypothermia and exsanguination.

### Sample collection and study design

Matings were assumed to occur at midnight, thus first detection of a vaginal plug was designated as 0.5 days post coitum, or E0.5, with subsequent days incremented accordingly (E1.5, E2.5 etc). Seven time points were used for whole embryo samples (E11.5, E12.5, E13.5, E14.5, E15.5, E16.5 and E18.5) with E13.5, E16.5 and E18.5 also used for head and forelimb samples. A minimum of three embryos were used for each time point, though where litter sizes permitted, greater numbers were collected (Whole embryos: E13.5 N=4, E18.5 N=5; Heads: E18.5 N=5, for a total of 24 whole embryo samples, 11 head samples and 9 forelimb samples –see Supplementary Table S1). Given potential variation in precise mating, ovulation and conception times, gestational ages should be considered approximate, particularly where different litters are pooled for a single timepoint (E13.5, E14.5, E18.5). At the appropriate gestational stage (all sample collections were performed between midday and 2pm), pregnant females were killed by cervical dislocation. Uterine horns were quickly removed and placed on ice (additional organs and muscles were collected from the adults for separate studies). After death, embryos were dissected from their uterine environments (including removal from amniotic sac) and either kept intact (whole embryos), or further dissected to isolate forelimbs (each sample used both forelimbs) and heads (remaining tissue from the body was also collected). All samples were snap-frozen in liquid nitrogen and stored at −80°C until use.

### RNA isolation and qPCR

Tissues (whole embryos, heads or forelimbs) were pulverised via dry-ice-cooled cell-crusher (Cellcrusher Ltd), and ~100-200mg frozen tissue powder (well mixed to ensure representative sampling) was used to prepare RNA. RNA was isolated using TRIzol (Invitrogen) as described previously [26, 37], with inclusion of an additional chloroform extraction (1:1) after the phase separation step and inclusion of 10μg glycogen during precipitation to maximise RNA yield. RNA yield and purity were assessed via nanodrop (ND1000) and samples with 260/230 ratios below 1.7 were subjected to a second precipitation. All cDNA was prepared using the RTnanoscript2 kit (Primerdesign), using 1600ng of RNA per reaction, with oligo dT and random nonamer priming. All reactions were subsequently diluted 1/20 with nuclease-free water to minimise downstream PCR inhibition. qPCR reactions were performed in duplicate or triplicate in 10μl volumes using 2μl diluted cDNA (~8ng cDNA per well assuming 1:1 conversion) in a CFX384 lightcycler using PrecisionPLUS SYBR green qPCR mastermix (Primerdesign), with a melt curve included as standard (Supplementary Figure S1). All Cq values were determined via linear regression. Primers to *ACTB, SDHA, GAPDH, HTATSF1, CDC40, RPL13A, CSNK2A2, AP3D1, CYC1, EIF4A, UBC, B2M* and *PAK1IP1* were taken from the geNorm and geNorm PLUS kits (Primerdesign): all give efficiencies of 95-105% and produce single amplicons. Sequences are proprietary but anchor nucleotides and context sequences can be provided on request. Primers to *18S* were those used previously [25], and those to *HPRT1* were the pan-species set (HPSF) validated by Valadan *et al* [48]. Primers targeting the 5’ terminus of dystrophin dp71 were designed using Primer3 (https://primer3.ut.ee/). All sequences are provided below:

18S F 5’-GTAACCCGTTGAACCCCATT-3’
18S R 5’-CCATCCAATCGGTAGTAGCG-3’
HPSF F 5’-GGACTAATTATGGACAGGACTG-3’
HPSF R 5’-GCTCTTCAGTCTGATAAAATCTAC-3’
Dp71 F 5’-GTGAAACCCTTACAACCATGAG-3’
Dp71 R 5’-CTTCTGGAGCCTTCTGAGC-3’

### Data analysis

All data was analysed using Microsoft Excel, using four different reference gene analysis programs: geNorm [33], deltaCt [36], BestKeeper [35] and Normfinder [34] (see supplementary material). BestKeeper and deltaCt methods used mean Cq values, while for geNorm and Normfinder Cq values were first linearised to relative quantities (RQ). All data were analysed within a single dataset (44 samples) or as embryos (24 samples), heads (11) or forelimbs (9) alone. Normfinder analysis further allows designation of user-defined groups: accordingly, data was analysed ungrouped as for the other packages above, or grouped as follows. Whole dataset, grouped by tissue type or age; whole embryos, grouped by age; heads grouped by age; forelimbs grouped by age.

To integrate the outputs of all programs, the per-gene geometric mean was generated from the scores of each specific comparison (whole dataset, embryos, heads, forelimbs). Bestkeeper ranks genes by coefficient of correlation (where low values represent poor correlation), while all other methods rank by stability (were low values represent high stability). Accordingly, values for Bestkeeper were inverted (1-value), with any negative correlation coefficients first set to zero (to give a subsequent score of 1). GeNorm suggests a best pair: for comparative purposes, each of the best pair were assigned the same score. Data from Normfinder was ungrouped (grouped analysis was not used in the integrated assessment).

### Statistical analysis

Statistical analysis was performed using Graphpad Prism 8.0 (Pearson correlations) or Microsoft Excel (coefficients of variation). Dataset coefficients of variation were determined for each timepoint individually, then summed to provide the overall variation.

## Results

### Raw Cq values

Raw Cq data (all genes, all samples) serves as a simple first-pass assessment of a reference gene panel. Most genes showed highly consistent expression across all samples (particularly *UBC*), however expression of *18S* was unexpectedly varied (Fig 2A): closer examination of the data suggested that levels of this gene varied not by age or tissue, but by cDNA synthesis batch, with one batch producing *18S* Cqs ~3 cycles later (corresponding to a roughly 10-fold reduction in template). Preparation of fresh cDNA for these samples corrected this discrepancy (Fig 2B). Given the comparatively consistent expression for other genes in our panel, we investigated further: restriction to *18S* signal only implied a defect specifically in random priming. Random priming is not required for cDNA synthesis *per se*, but is necessary for reverse transcription of ribosomal RNAs (which lack polyA tails) and more importantly, to capture 5’ sequence of longer mRNAs: deficient random priming would be expected to lower apparent expression of such mRNAs. To confirm this we measured expression of the dystrophin isoform dp71: this isoform is modestly expressed in the developing embryo, can only be distinguished from other dystrophin isoforms by primers targeting the unique first exon, and with a transcript ~5kb in length, capture of the 5’ terminus requires random priming. As shown (Fig 2C), measured Cq values for this isoform showed a very similar pattern of batch-specific variation which was similarly corrected following preparation of fresh cDNA from these samples.

**Fig 2:**
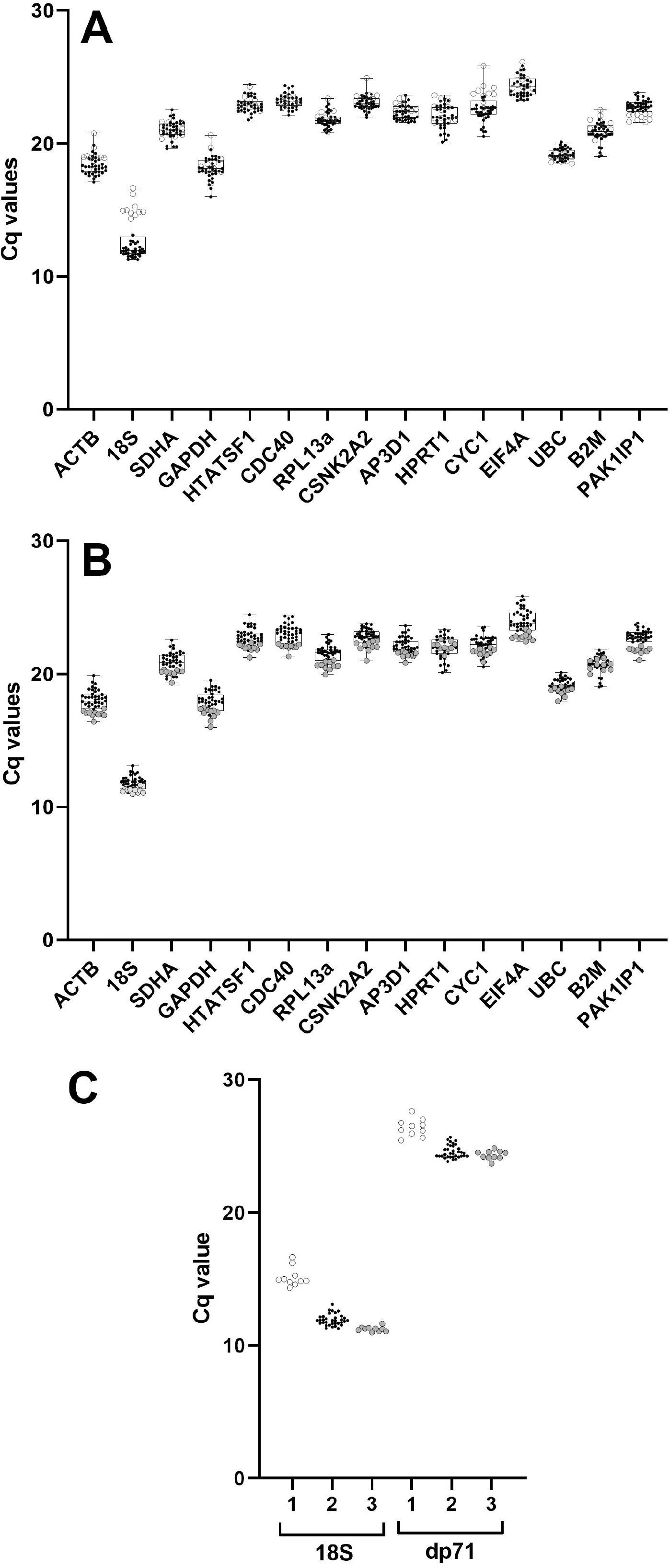
Raw Cq values and validation Box and whisker plots of sample Cq values for each candidate gene used in this study: points represent individual sample Cq values. (A) Initial raw Cq values revealed a marked variance in 18S expression (but not other genes), restricted to samples from a single cDNA synthesis batch (open circles), compared to samples from other cDNA synthesis batches (black points), indicative of impaired random priming. (B) Preparation of fresh cDNA for these samples corrected this discrepancy (filled circles), producing values consistent with other batches (black points). (C) qPCR for 18S and the dystrophin isoform dp71 using the initial poor samples (1), the other samples (2) and the fresh preparation (3) shows that impaired random priming prevents accurate quantification of long transcripts.

Assessment of our corrected dataset (Fig 2B) was largely in line with expected expression profiles, with known highly-abundant RNAs (*18S*, *ACTB*, *GAPDH*) showing the lowest mean Cq values. Interestingly, *EIF4A*, an RNA helicase that unwinds mRNA secondary structure for translational initiation, was the least abundant transcript in our panel. Given the fundamental role protein synthesis plays, and the high translational demands of a growing embryo, this finding was unexpected (though we note that even as the lowest expression gene in our panel, levels of this transcript were still comparatively high: mean Cq of ~23.9). Studies have suggested that initiation via the eIF4 protein complex involves a high degree of recycling at the 5’ UTR, with the same complex being reused multiple times in rapid succession: initiation complexes might be in lower overall demand than genes associated with metabolism or kinase signalling.

### geNorm analysis

The iterative geNorm algorithm ranks candidate genes by their mean pairwise variation M (where high M represents high variation) to determine the ‘best pair’, the pair of genes that correlate most closely across all samples. Low M values thus represent more suitable genes, but there is no fixed threshold: conventionally, M values <0.5 are considered acceptable as reference genes, while for samples with innately variable expression patterns (such as different tissues) values <1.0 may be accepted. GeNorm analysis (Fig 3, Table 2) revealed high overall stability: assessment of our dataset as a whole (all samples, Fig 3A) revealed 14 of our 15 candidate genes had acceptable M values (M<0.5). *CDC40* and *HTATSF1* were the ‘best pair’, though *RPL13A*, *PAK1IP1* and *ACTB* also performed well. *CYC1*, *GAPDH*, *B2M* and *HPRT1* all performed poorly, though only the latter failed to clear the M<0.5 threshold. We repeated the analysis using specific subsets of our data (embryos, heads or forelimbs only: Fig 3B, C and D respectively). Embryo samples alone gave results very similar to the dataset as a whole, both in ranking and magnitude of stability: *CDC40* and *HTATSF1* were again the best pair, with *RPL13A*, *PAK1IP1* and *ACTB* also scoring highly, while *CYC1*, *GAPDH*, *B2M* and *HPRT1* scored poorly. In heads alone, *CDC40* and *HTATSF1* were also the best pair, with *RPL13A* and *ACTB* being similarly high scoring (though here *GAPDH* also performed well, while *PAK1IP1* ranked poorly), but stability overall was greater in this tissue: all M values were <0.5, with 8 of the 15 showing M values <0.2. Greater overall stability was similarly revealed by analysis of forelimbs: a different best pair was obtained here (*ACTB* replacing *CDC40*), though *CDC40* (along with *RPL13a*) remained high scoring.

**Fig 3:**
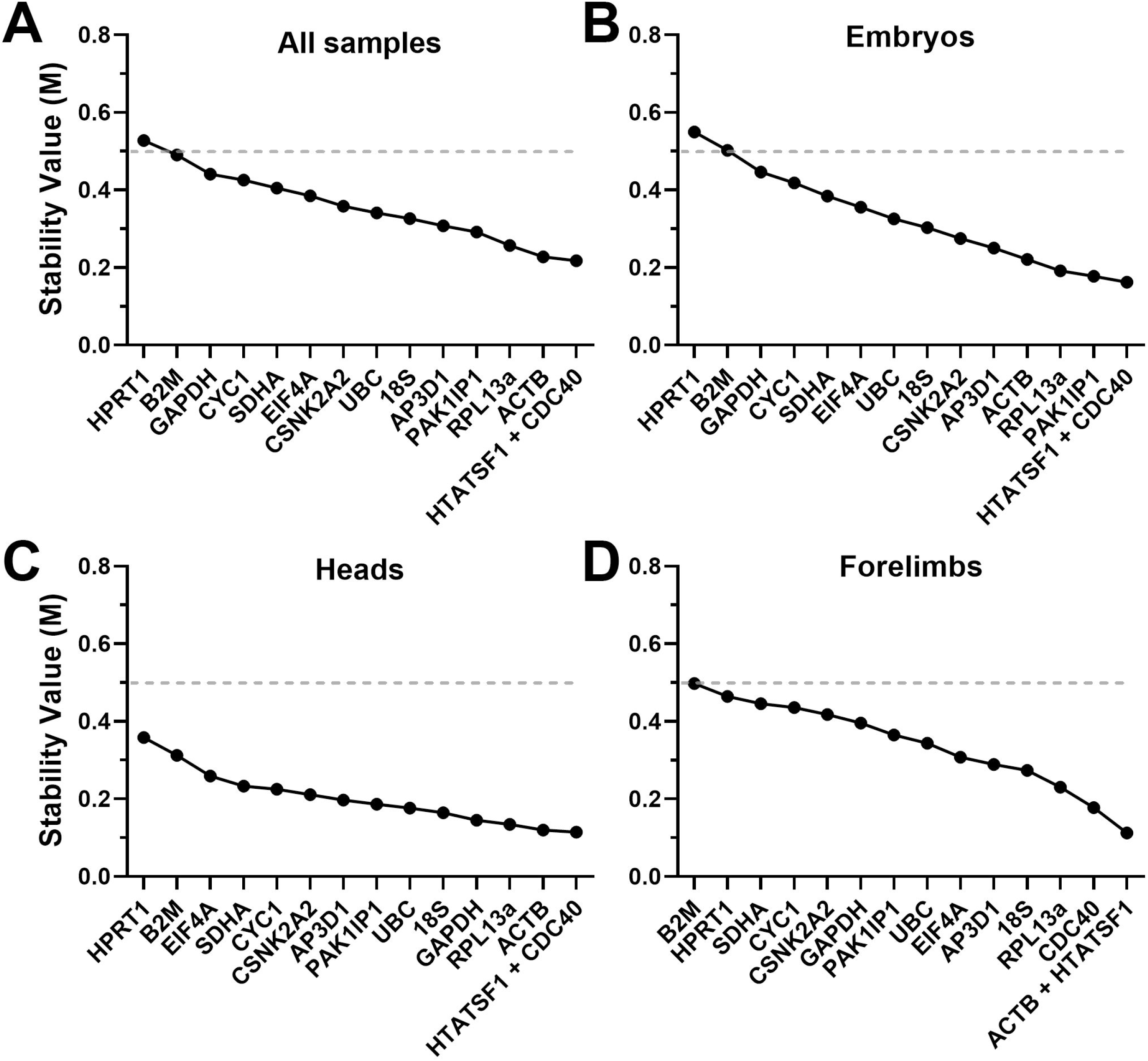
GeNorm rankings Rankings and scores of the fifteen candidate genes assessed by the geNorm algorithm, using the entire dataset (A), whole embryo samples only (B), heads only (C) or forelimbs only (D). High M values represent less stable genes. Genes with M values <0.5 (dashed line) are considered acceptable references.

**Table 2:**
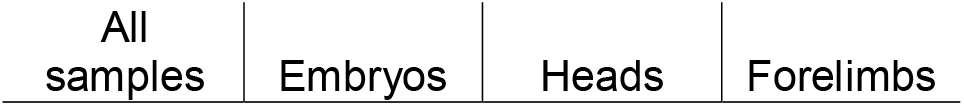
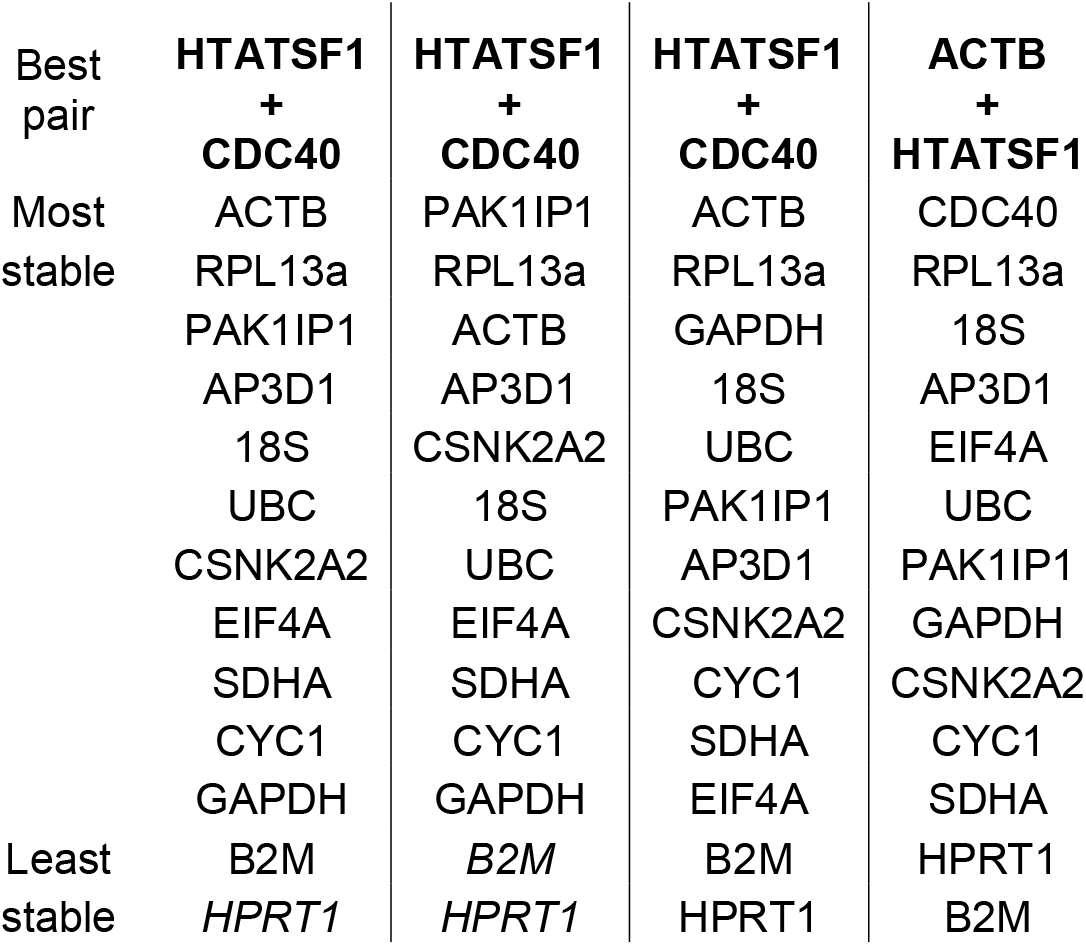
geNorm rankings GeNorm results for the entire dataset or tissue-specific subsets (as indicated), ranked from most stable to least stable. Genes with stability value M > 0.5 –poor scoring candidates- are indicated by italics.

| All samples | Embryos | Heads | Forelimbs |
| --- | --- | --- | --- |

|  |  |  |  |  |
| --- | --- | --- | --- | --- |
| Best pair | **HTATSF1**<br>+<br>**CDC40** | **HTATSF1**<br>+<br>**CDC40** | **HTATSF1**<br>+<br>**CDC40** | **ACTB**<br>+<br>**HTATSF1** |
| Most stable | ACTB | PAK1IP1 | ACTB | CDC40 |
|  | RPL13a | RPL13a | RPL13a | RPL13a |
|  | PAK1IP1 | ACTB | GAPDH | 18S |
|  | AP3D1 | AP3D1 | 18S | AP3D1 |
|  | 18S | CSNK2A2 | UBC | EIF4A |
|  | UBC | 18S | PAK1IP1 | UBC |
|  | CSNK2A2 | UBC | AP3D1 | PAK1IP1 |
|  | EIF4A | EIF4A | CSNK2A2 | GAPDH |
|  | SDHA | SDHA | CYC1 | CSNK2A2 |
|  | CYC1 | CYC1 | SDHA | CYC1 |
|  | GAPDH | GAPDH | EIF4A | SDHA |
| Least stable | B2M | *B2M* | B2M | HPRT1 |
|  | *HPRT1* | *HPRT1* | HPRT1 | B2M |

Determination of a best pair does not imply that only two reference genes should be used: geNorm analysis also therefore determines the change in pairwise variation elicited by increasing the number of reference genes. Typically, variation below 0.2 is considered acceptable, and while our data showed addition of a third reference gene lowered variation, the best pair alone was sufficient in all cases (Supplementary Fig S2).

### DeltaCt analysis

The DeltaCt (dCt) method ranks genes by the mean standard deviation of their pairwise Cq differences, with lower values indicating more stable references. Using this method (Fig 4, Table 3), comparable results were obtained from analysis of the dataset as a whole, or restricted to embryos only: sorting by mean deltaCt standard deviation revealed *HPRT1* and *B2M* were the worst performing candidates in both cases, by a considerable margin. *GAPDH*, *EIF4A* and *CYC1* also scored poorly, but modestly so: indeed, *HPRT1* and *B2M* aside, increases in apparent stability across the panel were slight, suggesting most of our panel represent valid reference genes (as suggested by geNorm, above). *AP3D1*, *PAK1IP1*, *CSNK2A2*, *RPL13A*, *UBC*, *CDC40* and *HTATSF1* all scored highly: despite slight differences in ranking between the whole dataset and embryos alone, all six genes had very similar scores (ranging from 0.42-0.5). Restricting analysis to the head alone (Fig 4C) produced a similar pattern of ranking, and increased apparent stability across the entire panel. With the exception of *HPRT1* and *B2M*, essentially every candidate gene represented a strong candidate for normalising gene expression in embryonic head tissues. *HPRT1* and *B2M* also ranked poorest in forelimbs alone (Fig 4D), but here *18S* and *EIF4A* were much higher scoring, while *ACTB* and *CDC40* were not, implying clear differences in expression patterns in these tissues (and a marked departure from geNorm, where *ACTB* formed part of the best pair).

**Fig 4:**
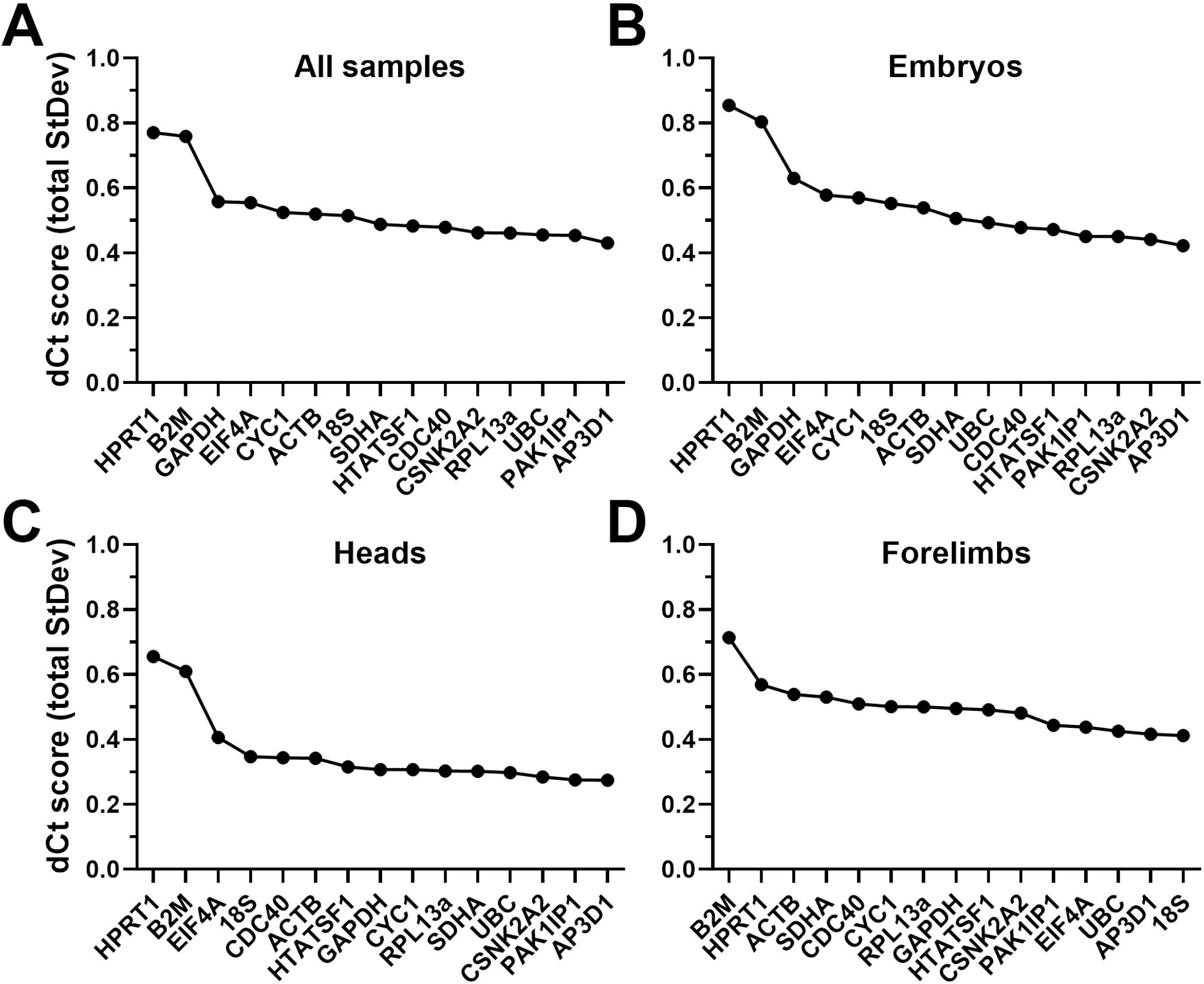
deltaCt rankings Rankings and scores of the fifteen candidate genes assessed by the deltaCt method, using the entire dataset (A), whole embryo samples only (B), heads only (C) or forelimbs only (D). High dCt scores represent less stable genes.

**Table 3:** DeltaCt rankings DeltaCt results for the entire dataset or tissue-specific subsets (as indicated), ranked from most stable to least stable. Genes with mean dCt standard deviations >0.6 –poor scoring candidates- are indicated by italics.

|  | All samples | Embryos | Heads | Forelimbs |
| --- | --- | --- | --- | --- |
| Most stable | AP3D1 | AP3D1 | AP3D1 | 18S |
|  | PAK1IP1 | CSNK2A2 | PAK1IP1 | AP3D1 |
|  | UBC | RPL13a | CSNK2A2 | UBC |
|  | RPL13a | PAK1IP1 | UBC | EIF4A |
|  | CSNK2A2 | HTATSF1 | SDHA | PAK1IP1 |
|  | CDC40 | CDC40 | RPL13a | CSNK2A2 |
|  | HTATSF1 | UBC | CYC1 | HTATSF1 |
|  | SDHA | SDHA | GAPDH | GAPDH |
|  | 18S | ACTB | HTATSF1 | RPL13a |
|  | ACTB | 18S | ACTB | CYC1 |
|  | CYC1 | CYC1 | CDC40 | CDC40 |
|  | EIF4A | EIF4A | 18S | SDHA |
|  | GAPDH | *GAPDH* | EIF4A | ACTB |
| Least stable | *B2M* | *B2M* | *B2M* | HPRT1 |
|  | *HPRT1* | *HPRT1* | *HPRT1* | *B2M* |

### Bestkeeper analysis

The Bestkeeper method averages the entire presented dataset to generate a consensus profile: the ‘Bestkeeper’, and then ranks individual genes by correlation with this. As shown above, analysis via geNorm and dCt tended to be more effective at highlighting poor genes from our panel than at identifying consistently strong candidates. Bestkeeper analysis added to this emerging pattern (Fig 5, Table 4): most candidates performed well, with *HPRT1* and *B2M* remaining the exceptions (exhibiting negative correlation with the Bestkeeper in the case of heads and forelimbs). Rankings of higher scoring genes disagreed with previous analyses, however: the gene *EIF4A* -ranked lower by both geNorm and dCt-correlated well with a Bestkeeper derived from the whole dataset (or from embryos only), while *ACTB*, *HTATSF1* and *CDC40* correlated less closely. Interestingly, *EIF4A* also correlated closely with a Bestkeeper derived from forelimb data only (Fig 5D), but not one derived from head samples (Fig 5C), while *CYC1* was high ranking in this latter tissue, but not in embryos, forelimbs or the dataset as a whole. *18S* was also closely correlated with the forelimb Bestkeeper (in agreement with dCt) but performed poorly in the other dataset groupings.

**Fig 5:**
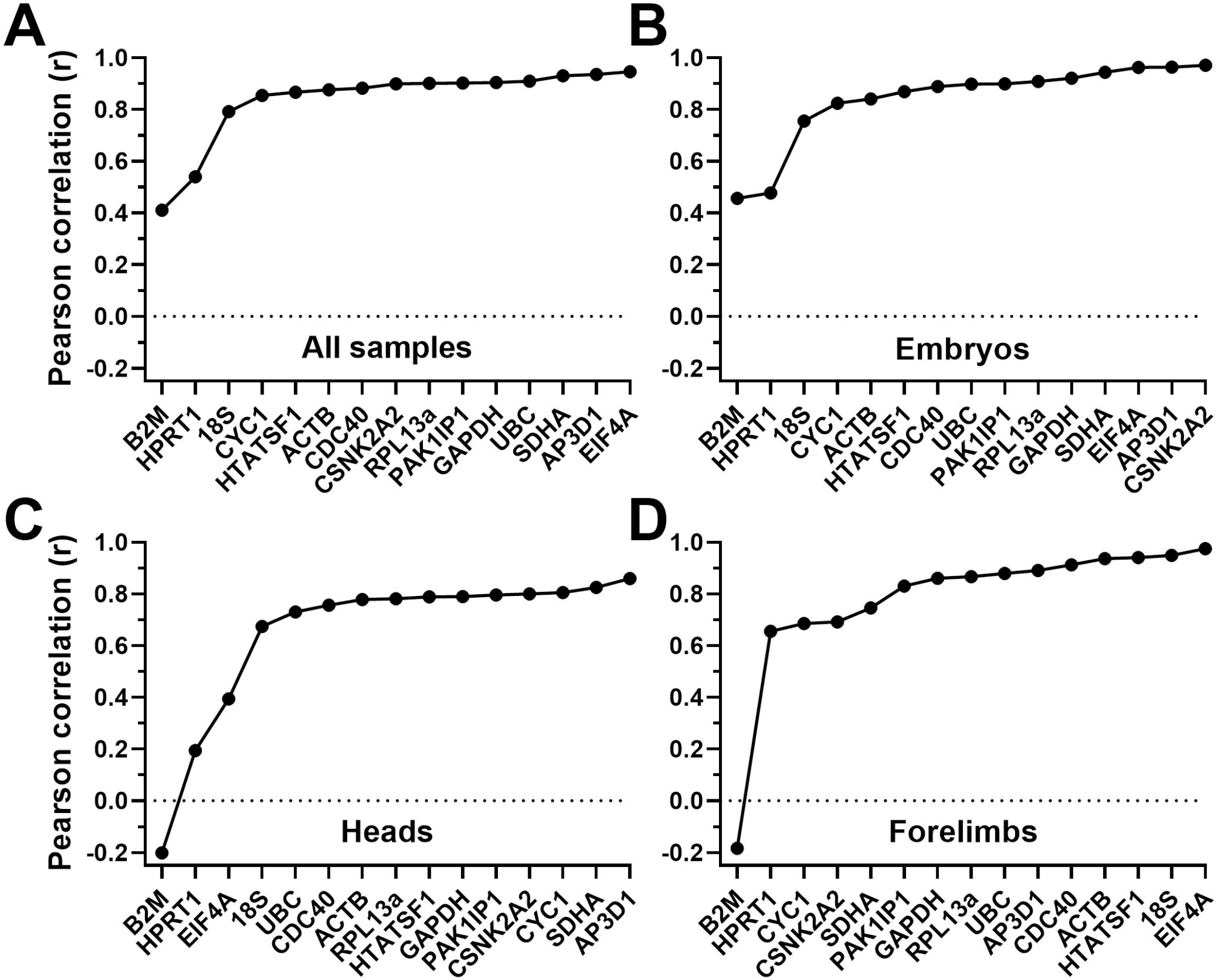
Bestkeeper rankings Rankings and scores of the fifteen candidate genes assessed by the Bestkeeper method, using the entire dataset (A), whole embryo samples only (B), heads only (C) or forelimbs only (D). Genes are ranked by their individual Pearson correlation coefficient (r) to the bestkeeper derived from all candidate genes: low r values correspond to less stable genes. Dashed line: r value of zero.

**Table 4:**
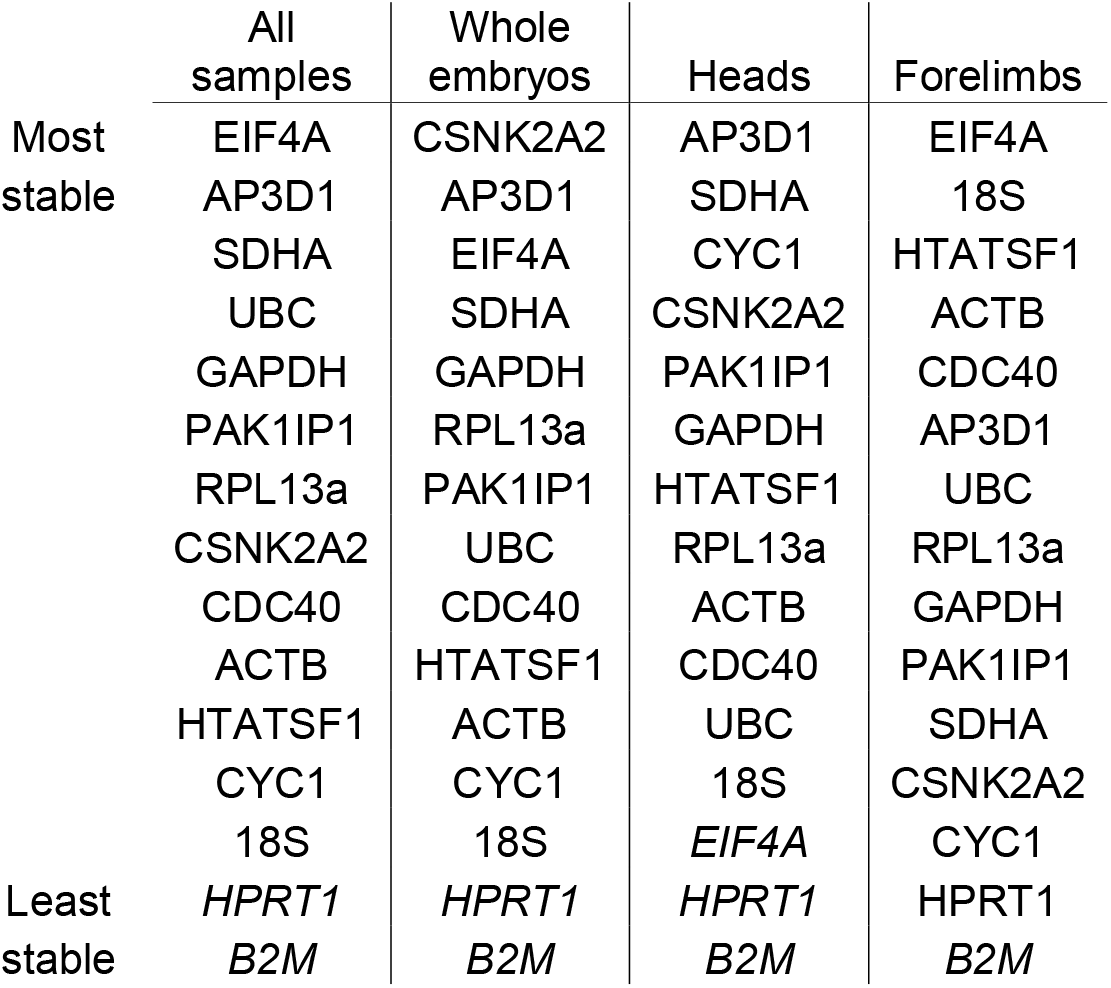
Bestkeeper rankings Bestkeeper results for the entire dataset or tissue-specific subsets (as indicated), ranked by Pearson correlation coefficient (r) with the Bestkeeper, from most stable to least stable. Genes with r <0.6 –poor scoring candidates- are indicated by italics.

### Normfinder analysis

Unlike the other three algorithms, Normfinder does not make pairwise comparisons, but instead assesses each gene for individual stability across the dataset. This method also allows two different analysis approaches: grouped and ungrouped. Ungrouped analysis considers the dataset without distinction between samples, while grouped analysis allows discrete subcategories to be defined within the dataset. Accordingly, we used ungrouped analysis to assess our combined dataset and tissue-restricted subsets (Fig 6, Table 5), with further grouped analysis to introduce age and tissue-type as subcategories within those datasets where appropriate. In some respects, ungrouped analysis agreed with the above methods (particularly dCt): *B2M* and *HPRT1* were consistently ranked last (by a considerable margin), while many of the remaining genes had comparable high scores. *EIF4A* ranked lower here in all tissues but forelimbs, while *CNSK2A2*, *PAK1IP1* and especially *AP3D1* all typically performed well (in agreement with dCt and geNorm, but in contrast to Bestkeeper). *18S* was again ranked highest in forelimbs alone, but *ACTB* fared very poorly (in agreement with dCt and Bestkeeper, but not geNorm)

**Fig 6:**
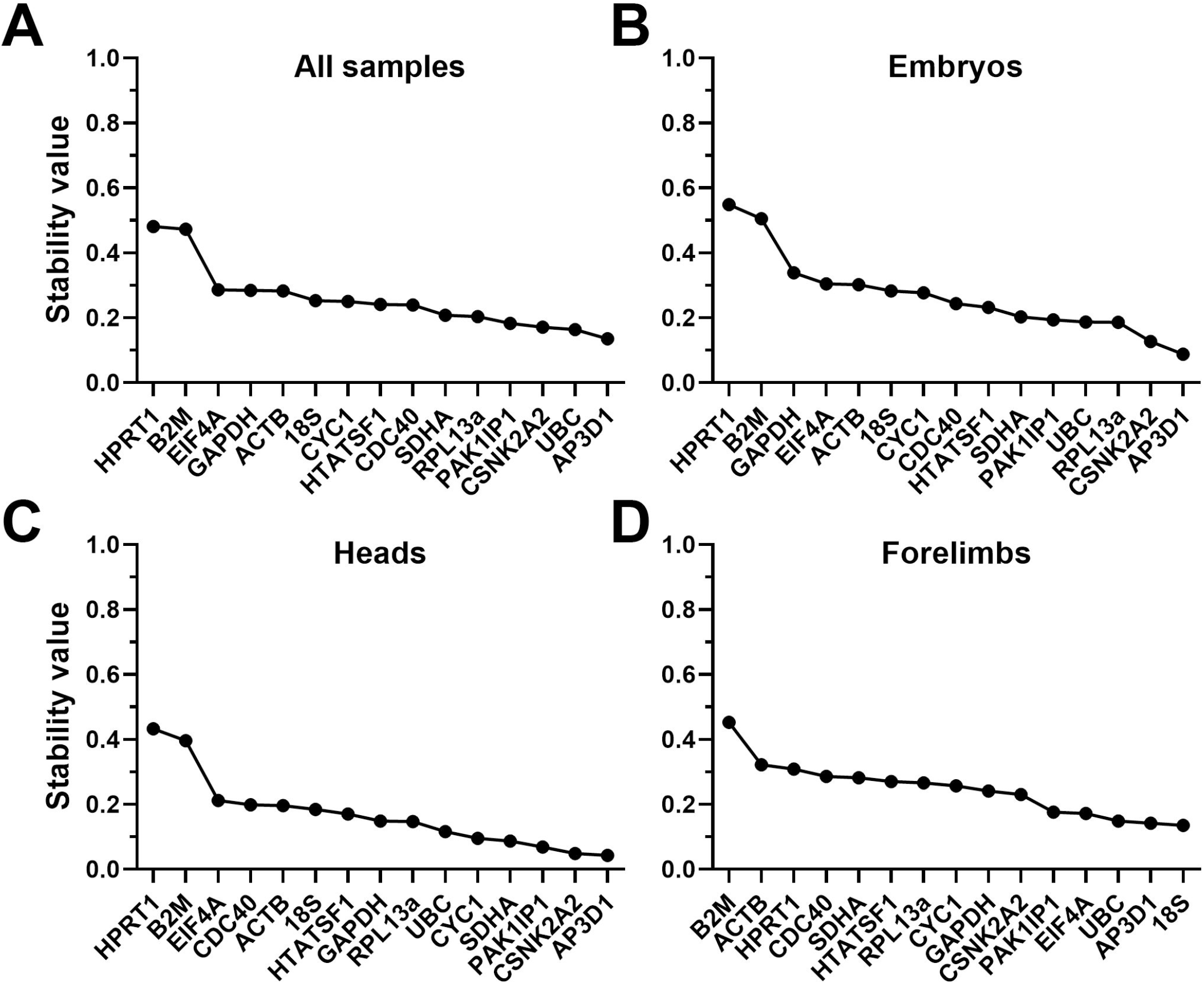
Normfinder rankings (ungrouped) Ungrouped rankings and scores of the fifteen candidate genes assessed by the Normfinder algorithm, using the entire dataset (A), whole embryo samples only (B), heads only (C) or forelimbs only (D). High stability values correspond to less stable genes.

**Table 5:**
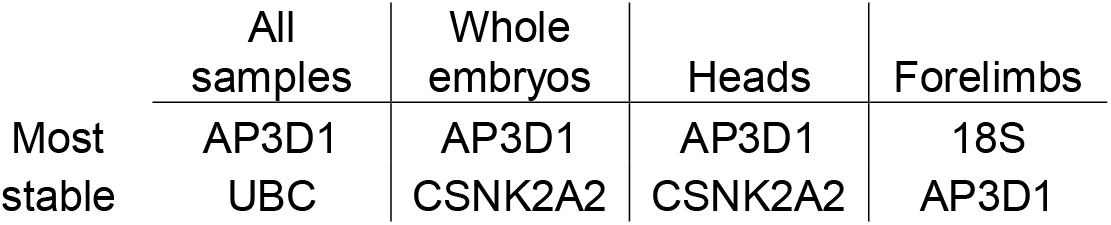
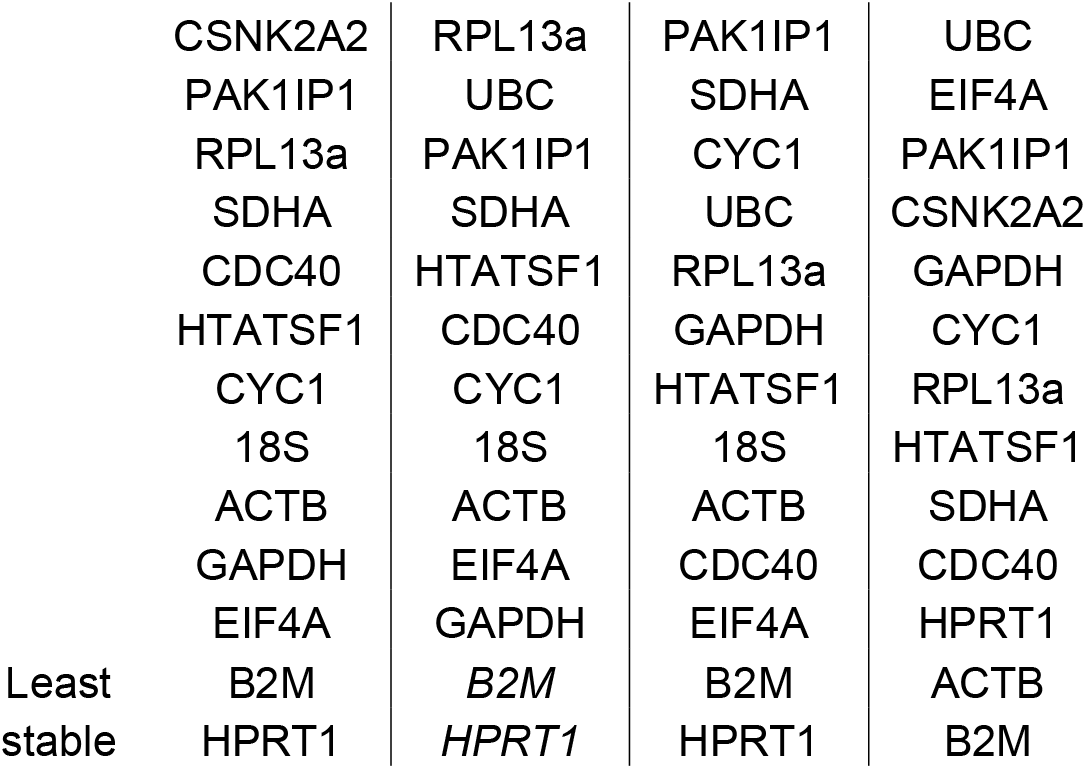
Normfinder rankings (ungrouped) Ungrouped Normfinder results for the entire dataset or tissue-specific subsets (as indicated), ranked from most stable to least stable. Genes with stability values >0.5 –poor scoring candidates- are indicated by italics.

|  | All samples | Whole embryos | Heads | Forelimbs |
| --- | --- | --- | --- | --- |
| Most stable | AP3D1 | AP3D1 | AP3D1 | 18S |
|  | UBC | CSNK2A2 | CSNK2A2 | AP3D1 |
|  | CSNK2A2 | RPL13a | PAK1IP1 | UBC |
|  | PAK1IP1 | UBC | SDHA | EIF4A |
|  | RPL13a | PAK1IP1 | CYC1 | PAK1IP1 |
|  | SDHA | SDHA | UBC | CSNK2A2 |
|  | CDC40 | HTATSF1 | RPL13a | GAPDH |
|  | HTATSF1 | CDC40 | GAPDH | CYC1 |
|  | CYC1 | CYC1 | HTATSF1 | RPL13a |
|  | 18S | 18S | 18S | HTATSF1 |
|  | ACTB | ACTB | ACTB | SDHA |
|  | GAPDH | EIF4A | CDC40 | CDC40 |
|  | EIF4A | GAPDH | EIF4A | HPRT1 |
| Least | B2M | *B2M* | B2M | ACTB |
| stable | HPRT1 | *HPRT1* | HPRT1 | B2M |

Grouped analysis of our datasets (adding in an age or tissue component to the combined data, and an age component to tissue-specific subsets) differed from ungrouped in several respects (Fig 7, Table 6), suggesting interesting nuances of expression stability within our datasets. When the entire dataset was grouped by tissue (embryo, head, forelimb) no gene had a stability value greater than 0.2, i.e. stability across the panel was substantially greater overall than for the ungrouped dataset, or the same data grouped by age. This suggests tissue-specific differences in gene expression might be mild (and certainly milder than age-specific changes). *HPRT1*, *B2M*, *EIF4A* and *18S* were the lowest ranked genes in both tissue and age-specific grouping, and indeed *HPRT1* and *B2M* scored poorly in all rankings, with the marked exception of embryos grouped by age (Fig 7C). Here *CYC1*, *PAK1IP1* and *18S* performed worse (exceptionally so, in the case of *18S*). Grouping embryos by age moreover revealed much lower overall stability for the panel than all other grouped analyses: these findings imply global age-related changes are of greatest magnitude, and that *CYC1*, *PAK1IP1* and *18S* in particular might exhibit consistent age-associated changes. Grouped analysis additionally generates a best pair: two genes that may vary between groups, but in opposite directions (given greater overall stability than any individual gene, when combined). In most cases, this best pair did not feature the most stable genes, but as the majority of genes performed well under the majority of comparisons, this was not unexpected (and the stability advantages offered by this best pair were in most cases only modest improvements over the highest scoring candidates alone).

**Fig 7:**
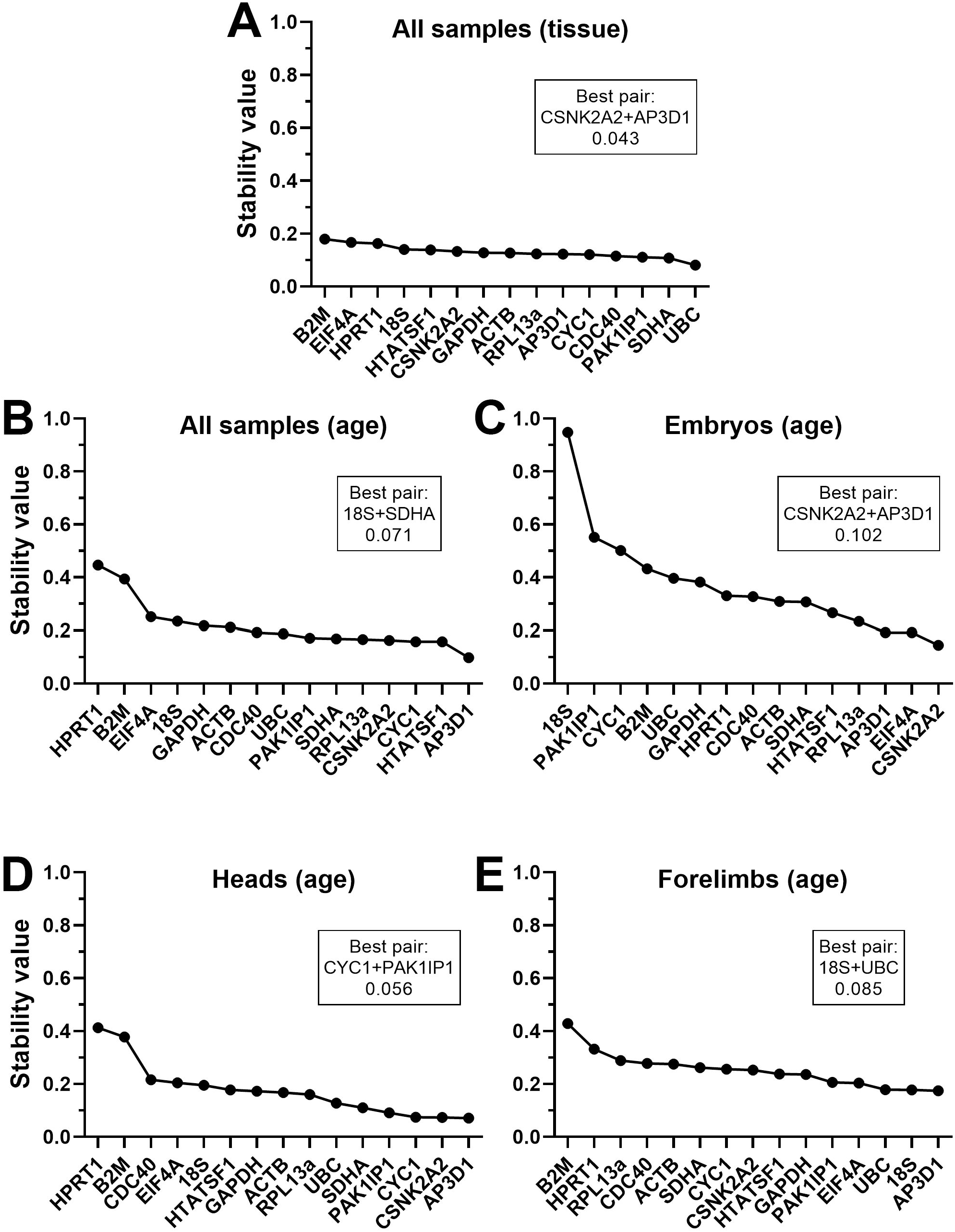
Normfinder rankings (grouped) Grouped rankings and scores of the fifteen candidate genes assessed by the Normfinder algorithm, using the entire dataset grouped by tissue (A) or by age (B), or whole embryo samples only grouped by age (C), heads grouped by age (D) or forelimbs grouped by age (E). High stability values correspond to less stable genes. The best pair of genes (which may not be the highest scoring individually) and the stability of that pair are indicated (boxes).

**Table 6:**
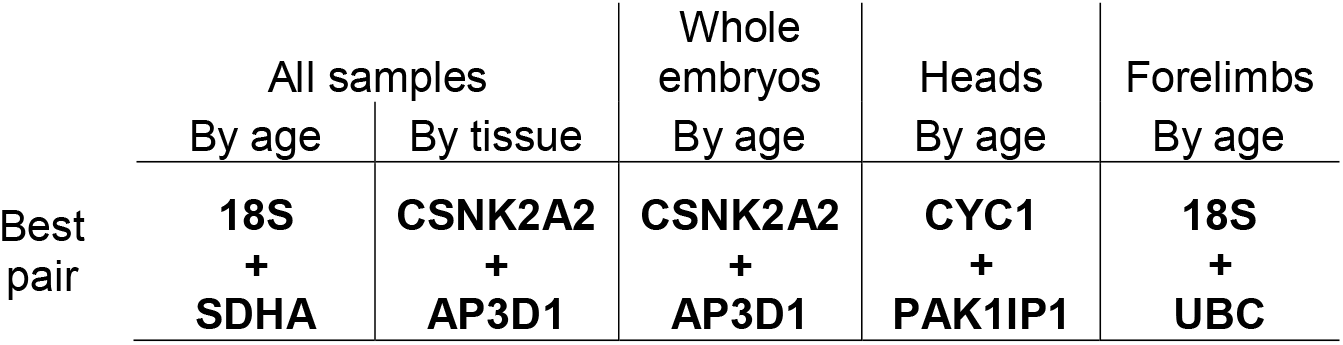
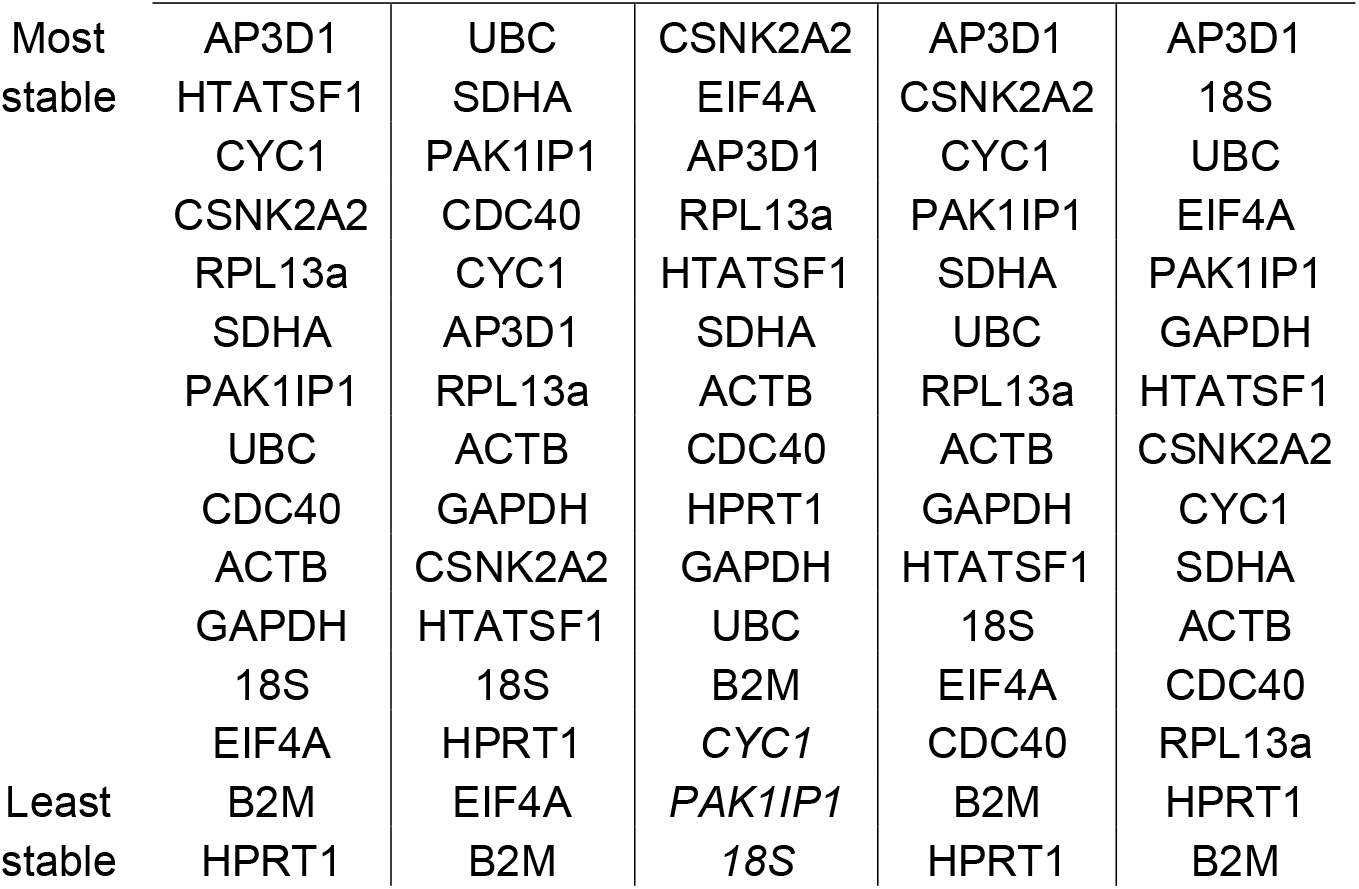
Normfinder rankings (grouped) Grouped Normfinder results for the entire dataset or tissue-specific subsets, grouped internally by age or tissue (as indicated), ranked from most stable to least stable. The best pair of genes are indicated separately (bold). Genes with stability values >0.5 –poor scoring candidates- are indicated by italics.

### Summary and Validation

Despite the differences in highest scoring candidates between algorithms, all four analysis methods tended to suggest that many of our candidate genes represented stable references. In most cases, the highest-ranking candidate was only fractionally more stable than candidates ranked seventh or eighth. Conversely, at the other end of the stability spectrum all algorithms were in much greater agreement. Regardless of method used or tissue assessed, *HPRT1* and *B2M* were near-universally ranked last, and often by a substantial margin, implying strongly that these genes are inappropriate choices. To illustrate this more clearly, we integrated our collected data in a manner similar to the RefFinder method developed by Xie *et al* [49], taking the geometric mean of all algorithm scores, either for the entire dataset, or for embryos, heads and forelimbs alone (see methods). When combined in this fashion (Fig 8, Table 7), the patterns and rankings are remarkably similar for all sets except forelimbs alone. In embryos, heads, or all samples together, *AP3D1* is the strongest candidate, while *RPL13A*, *PAK1IP1* and *CSNK2A2* are also highly ranked. In forelimbs, *EIF4A*, *18S*, *ACTB* and *HTATSF1* are the strongest candidates, however *AP3D1*, *RPL13A* and *PAK1IP1* also score highly in this tissue (while *CSNK2A2* does not). Taken together, these data suggest that *AP3D1*, *PAK1IP1* and *RPL13A* might represent a universally suitable panel.

**Fig 8:**
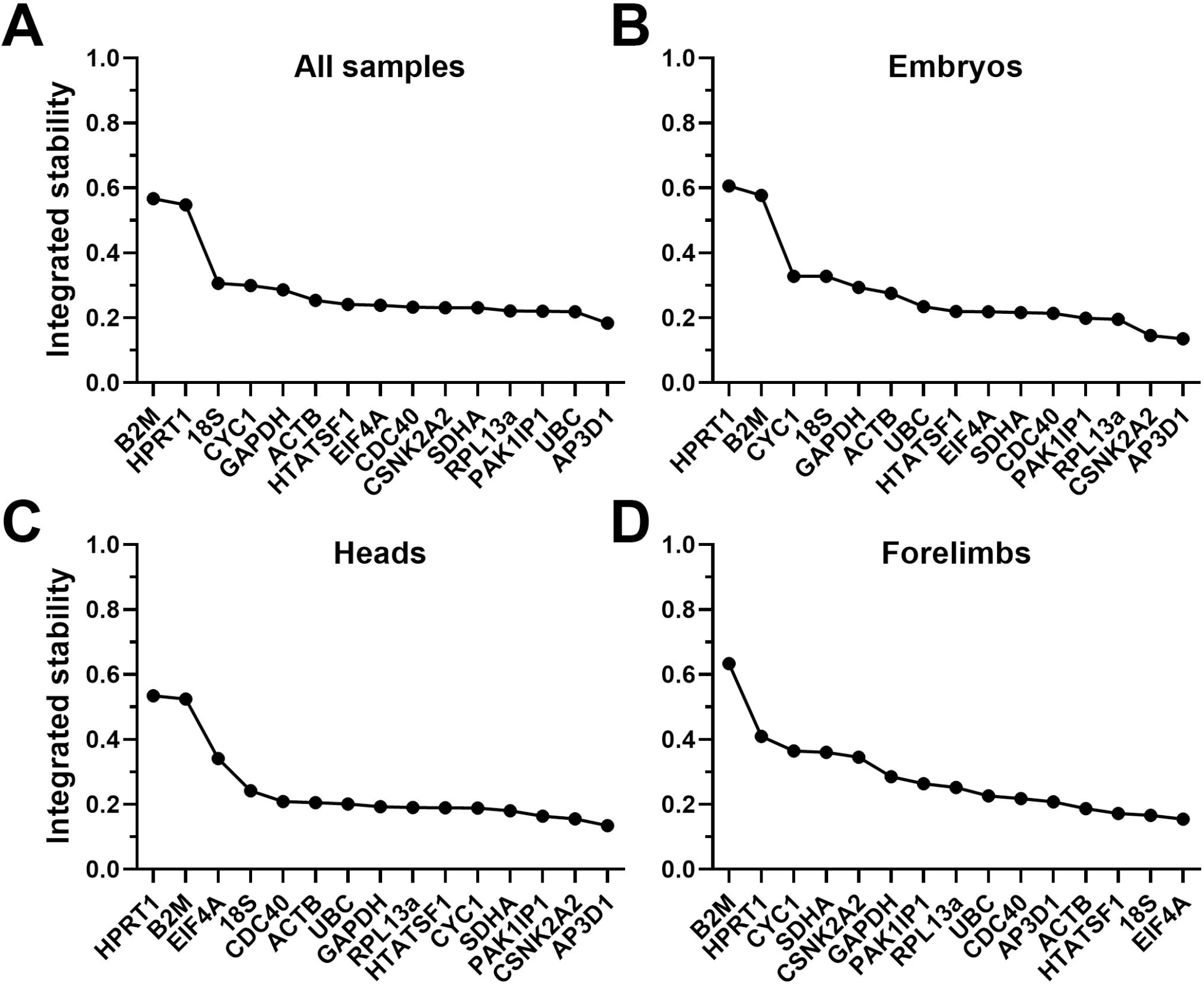
Integrated rankings of the four analysis methods Geometric mean scores of all fifteen candidate genes from all four algorithms (see methods), assessed as either a complete dataset (A), whole embryos only (B), heads (C) or forelimbs (D). High integrated stability values correspond to less stable genes.

**Table 7:** Integrated ranking Reference genes ranked by geometric mean of geNorm, dCt, Bestkeeper and Normfinder scores. Genes with stability values >0.5 –poor scoring candidates- are indicated by italics.

|  | All<br>samples | Whole<br>embryos | Heads | Forelimbs |
| --- | --- | --- | --- | --- |
| Most | AP3D1 | AP3D1 | AP3D1 | EIF4A |
| stable | UBC | CSNK2A2 | CSNK2A2 | 18S |
|  | PAK1IP1 | RPL13a | PAK1IP1 | HTATSF1 |
|  | RPL13a | PAK1IP1 | SDHA | ACTB |
|  | SDHA | CDC40 | CYC1 | AP3D1 |
|  | CSNK2A2 | SDHA | HTATSF1 | CDC40 |
|  | CDC40 | EIF4A | RPL13a | UBC |
|  | EIF4A | HTATSF1 | GAPDH | RPL13a |
|  | HTATSF1 | UBC | UBC | PAK1IP1 |
|  | ACTB | ACTB | ACTB | GAPDH |
|  | GAPDH | GAPDH | CDC40 | CSNK2A2 |
|  | CYC1 | 18S | 18S | SDHA |
|  | 18S | CYC1 | EIF4A | CYC1 |
| Least | *HPRT1* | *B2M* | *B2M* | *HPRT1* |
| stable | *B2M* | *HPRT1* | *HPRT1* | *B2M* |

For validation, we employed an approach we have used previously [26, 37]: using our high scoring candidates (*AP3D1*, *RPL13A*, *PAK1IP1*) to normalise our lowest scoring candidates (*HPRT1* and *B2M*). Raw data for these genes suggested modest expression with a dramatic increase at E18.5, however following normalisation both genes show a progressive increase in expression with increasing age, whether in whole embryos, heads, or limbs (Fig 9). Normalisation would also be expected to reduce the overall coefficient of variation (CoV), and as shown this was indeed the case.

**Fig 9:**
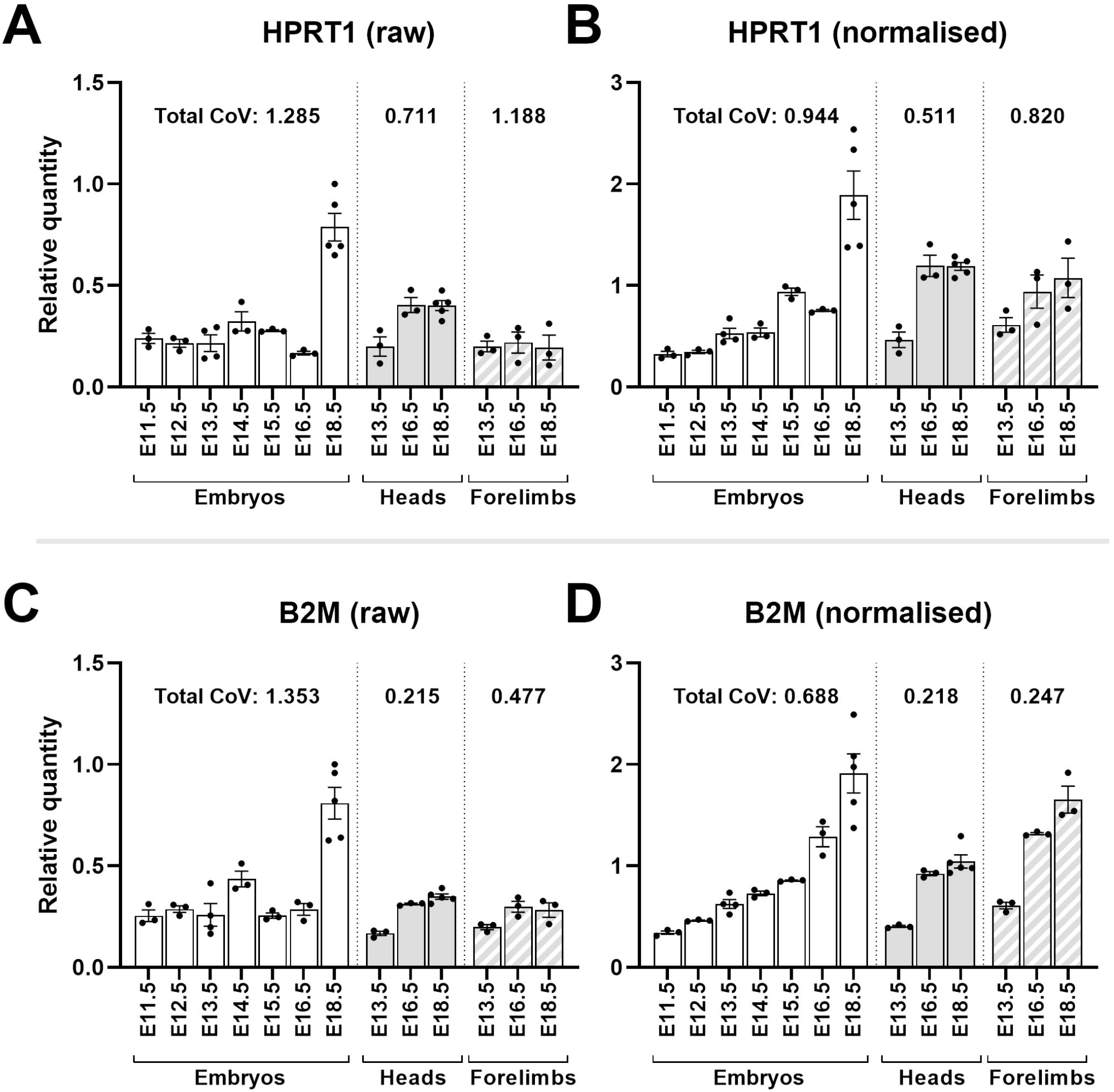
Normalisation reveals HPRT1 and B2M are developmentally upregulated Mean raw RQ values for HPRT1 (A) and B2M (C) suggest modest expression at earlier stages with marked increases at E18.5. Normalisation with AP3D1, RPL13A and PAK1IP1 (B and D) lowers total coefficient of variation (CoV) and shows both genes are upregulated with increasing gestational age in whole embryos, heads and forelimbs (as indicated). Total CoV values were obtained by summing the individual CoVs per time-point (see methods). Points represent individual RQ values (arbitrary units).

This validation method can be extended: the approaches used by each algorithm are subtly different, and thus information can be gleaned by examining cases where candidate rankings disagree. GeNorm, a method that ranks genes by pairwise variation, suggested *HTATSF1* and *CDC40* as the best pair near-unanimously, while these two genes performed only modestly under most other assessments. The implication is that these genes share a very similar pattern of expression, but in a manner unstable across samples. To investigate this directly, we used our high scoring candidates to normalise expression of *HTATSF1* and *CDC40*. Raw data was indeed highly variable between samples, and between tissues and ages, yet this variability was well-matched between the two genes, confirming their geNorm score (Fig 10). Following normalisation (which again substantially lowered CoV), this variability resolved into a mild, but remarkably consistent, decrease in expression with increasing age, in all tissues. Normalisation of *CYC1*, *UBC* or *EIF4A* (Supplementary Fig S3) gave similar results: *CYC1*, which typically scored highly in heads but low elsewhere, was revealed to be very consistently expressed in the former tissue while exhibiting age-associated increases in embryos and forelimbs (Supplementary Fig S3A and B); *UBC*, which was ranked modestly overall, exhibited a similar age-associated increase in expression in all tissues except the head (Supplementary Fig S3C and D); *EIF4A*, which scored particularly highly in forelimbs, was indeed stable in this tissue, while showing age-specific increases in heads and highly variable behaviour in whole embryos (Supplementary Fig S3E and F). Given *18S* and *EIF4A* were the highest scoring candidates in forelimbs, we further compared normalisation using these two genes with that obtained using the three genes above. Normalised forelimb expression of *B2M*, *HPRT1*, *CDC40* and *HTATSF1* was comparable with either reference gene combination, though overall CoV values were lower using *AP3D1*, *RPL13A* and *PAK1IP1* (Supplementary Fig S4).

**Fig 10:**
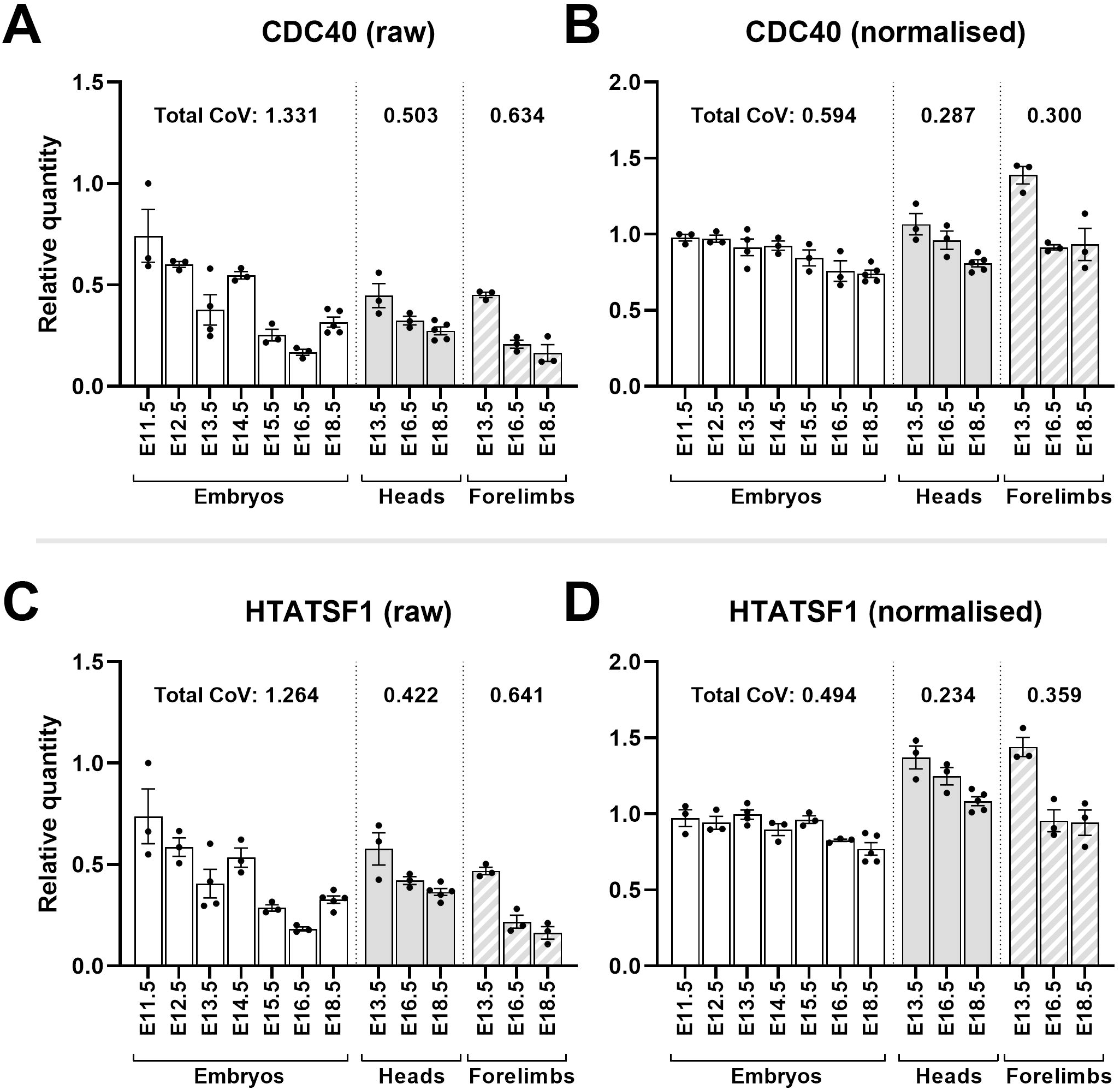
Normalisation reveals CDC40 and HTATSF1 are developmentally downregulated Mean raw RQ values for CDC40 (A) and HTATSF1 (C) suggest closely-matched but variable expression. Normalisation with AP3D1, RPL13A and PAK1IP1 (B and D) lowers total coefficient of variation (CoV) and shows both genes are modestly downregulated with increasing gestational age in whole embryos, and more markedly downregulated in heads and forelimbs (as indicated). Total CoV values were obtained by summing the individual CoVs per time-point (see methods). Points represent individual RQ values (arbitrary units).

Finally, use of all three reference genes might not be necessary: while the MIQE guidelines suggest use of two reference genes at a minimum [24], our geNorm analysis suggested no substantial benefit in increasing number of genes from 2 to 3 (or indeed from 3 to 4).

Accordingly, we compared a normalisation factor (NF) using all three genes to one using *AP3D1* with either *RPL13A* or *PAK1IP1* alone: the three gene NF was essentially identical to either 2-gene NF (gradients of 1.00 and 0.978 respectively, both with Pearson correlations of 0.99 -Supplementary Fig S5), suggesting that two genes (*AP3D1* plus one other) are indeed sufficient.

## Discussion

The later stages of embryonic development involve many significant changes: substantial organogenesis occurs over this period, as does formation of skeletal muscle and the skeleton itself. Within the head, the brain matures from a comparatively simple tube to an intricately subdivided organ complete with discrete ventricles and regional specialisation, the teeth develop from buds to near-maturity, while the eye progresses from the simple optic cup stage to a defined globe complete with lens and iris, hidden beneath the fused eyelids. Normalisation of gene expression throughout this period might well be expected to be challenging, however our data unexpectedly suggests otherwise. For our dataset, whether assessed in its entirety, or restricted to whole embryos, heads or forelimbs alone, all four analysis methods suggested that a substantial number of our candidate genes would serve as suitable reference genes: with the exception of *HPRT1* and *B2M*, essentially any gene from our panel represents an adequate reference. Given our panel consists of genes specifically selected for their reported stability, this is perhaps more reassuring than not, however we note that use of a near-identical panel of genes in skeletal muscle revealed markedly lower overall stability (for geNorm in particular, only *ACTB* and *RPL13a* gave M values <0.5) [37]. Our findings here imply that the developing embryo might be more transcriptionally consistent than different mature skeletal muscles. Even with most genes scoring highly there were differences, however, both between algorithms and between tissues, but the combination of *AP3D1*, *RPL13A* and *PAK1IP1* (or indeed *AP3D1* plus one of the others) emerged as consistently stable overall, and thus represent universal reference genes for the developing mouse embryo. We note that *18S* and *EIF4A* apparently represent better candidates in forelimbs specifically, but only fractionally so: studies addressing developmental changes in limbs alone might technically be better served by these two, but this would necessarily sacrifice the utility advantages offered by a universal panel (and *AP3D1*, *RPL13A* and *PAK1IP1* perform comparably to *18S*/*EIF4A* in forelimbs - Supplementary Fig S4).

AP3D1 codes for a component of the adaptor complex, which mediates non-clathrin-coated vesicle trafficking: we have previously found this gene to be stable throughout myogenic differentiation in culture [25] and in mature mouse skeletal muscle (healthy and dystrophic) [37], and our data here support the possibility that this gene might exhibit high overall stability in mouse. *RPL13A* codes for a protein component of the small ribosomal subunit: ribosomal proteins are ubiquitous, and some have proposed that such ubiquity renders them generally suitable as references [50], though this remains contentious [51]. As with *AP3D1*, we have previously found this gene to be stable in mature mouse skeletal muscle, and indeed also suitable as a reference in canine skeletal muscle (again, both healthy and dystrophic [26]). *PAK1IP1* encodes a negative regulator of PAK1 kinase, and its presence in our top scoring candidates represents something of a surprise: this gene was included in our candidate panels for both myogenic cell cultures and mature mouse muscle [25, 37], and in those scenarios its performance was near-uniformly mediocre. Here this gene was ranked lower even than *HPRT1* and *B2M* in Normfinder analysis of embryos grouped by age, but outside of this specific scenario the gene performed well, and as shown (Supplementary Fig S5), *RPL13A* and *PAK1IP1* are essentially interchangeable when used in combination with *AP3D1*.

Our validation process confirmed the utility of these genes, and also revealed insight into changes in gene expression during development: normalised expression of *HPRT1* and *B2M* (our lowest scoring genes) exhibited clear age-associated upregulation (Fig 9). *B2M* (beta 2 microglobulin) is a component of the MHC class I complex, and while this complex is present even at the earliest stages of development, studies in the rat have shown marked increases in expression in specific tissues such as skin, lung and inter-organ connective tissue [52]. Many of these tissues are only emerging at the earliest stages of our sample set (E11.5-E13.5) and indeed the skin barrier is complete only toward the end of development (E16.5-E18.5) [18]. A developmentally associated upregulation in *B2M* is consistent with these changes. *HPRT1* encodes Hypoxanthine Phosphoribosyltransferase 1, a central component of the purine salvage pathway: this gene is stably expressed under many circumstances (we have reported this gene to be stable in healthy and dystrophic canine muscle [26]). Mutations in this gene however cause the neurodevelopmental condition, Lesch-Nyhan disease, and the gene itself plays a key role in glial/neuronal fate choice [53], implying that expression is developmentally regulated. Our data supports this, and indeed suggests that expression of this gene increases earlier in the head than within the embryo as a whole (Fig 9B, expression at E16.5). Conversely, both *CDC40* and *HTATSF1* displayed age-correlated downregulation following normalisation (Fig 10): this decrease was modest (insufficient to exclude them as acceptable, though not optimal, references) and moreover was sufficiently consistent between the two genes as to flag them as the best pair under geNorm analysis. This finding serves as a prominent reminder of the potential risks in relying on a single reference gene analysis method, but also indicates these genes might be co-ordinately regulated. *HTATSF1* encodes HIV-TAT specific factor 1, an RNA binding protein. Recent studies in embryonic stem cells implicate this factor in both regulation of ribosomal RNA processing and splicing of mRNA for ribosomal proteins, influencing protein synthesis and cell differentiation as a consequence [54]. *CDC40* (cell division cycle 40, also known as *PRP17*, or pre-mRNA processing factor 17) encodes a protein component of the spliceosome, and is also involved in progression through the cell cycle. Given our data, it seems likely that a shared involvement in mRNA splicing underpins the remarkably well-matched expression of these two genes, with requirements for spliceosomal components decreasing gradually with developmental progress.

Our dataset also revealed a marked age-related increase in *CYC1* expression in all tissues except the head (Supplementary Fig S3B). This gene encodes the mitochondrial electron transfer chain component cytochrome C, and our findings are thus best explained by developmental mitochondrial biogenesis. The most likely source of such marked increases in mitochondrial content is skeletal muscle: E11.5 represents the approximate midpoint of primary myogenesis, in which small numbers of myofibres are laid down to serve as scaffolds for subsequent musculature, while the period from E14.5-E18.5 spans secondary myogenesis, a phase of more substantial muscle synthesis [10]. This latter phase might well be associated with a dramatic increase in mitochondrial content, both in the embryo as a whole but especially in the forelimb where skeletal muscle comprises a greater fraction of the total tissue. The head is comparatively muscle-poor by contrast, and craniofacial muscles are moreover derived from a different progenitor pool which need not mature along identical timescales [55]. The brain is a highly energetic tissue, however neurodevelopmental changes over this period are predominantly structural/differentiation rather than bulk increases in tissue mass. In contrast, *EIF4A* showed the reverse trend (Supplementary Fig S3F), increasing in heads but not forelimbs or embryos as a whole. This gene codes for an RNA helicase that unwinds 5’ secondary structure for the translational preinitiation complex, and increases in expression in the head over this period might reflect the demands of the diverse transcription/translation programs necessary for brain maturation.

Finally, our experience with *18S* merits additional mention. Ribosomal RNAs comprise the vast bulk of total RNA (~80-85%), but the absence of polyA tails necessitates random priming for conversion to cDNA. Oligo dT does not contribute to *18S* cDNA, thus this ribosomal target effectively serves as an internal control for efficiency of random priming. As we show here, batch-dependent variance in *18S* specifically (i.e. other genes remain comparable) is highly indicative of a failure in this specific step, perhaps due to degradation of random primer stocks. Under many circumstances, this might be of little consequence: while the processivity of reverse transcriptase is such that even highly optimised enzymes seldom incorporate more than 1500 nucleotides in a single binding event [56], the majority of mRNAs are of modest (~3kb) length [57] and can typically be reverse-transcribed in their entirety from initial oligo dT priming alone. 5’ sequence of longer transcripts (5kb+) will however be increasingly underrepresented or even absent under such conditions, and random priming is thus essential for qPCR studies investigating such transcripts (such as the early embryonic dystrophin isoform dp71). Critically, a failure in random priming will not be immediately apparent if all reference genes used can be successfully reverse-transcribed via oligo dT: our data suggests that for studies using random priming in combination with oligo dT, particularly those investigating expression of long genes, measurement of *18S* could represent a prudent quality check regardless of its efficacy as a reference.

## Conclusion

We have investigated potential reference genes for normalising quantitative PCR expression data in the developing mouse embryo from E11.5 to E18.5. We investigated expression both in whole embryos, or heads or forelimbs in isolation. Our data suggests that normalisation of expression over this period is not only possible, but that many reference genes are acceptable, though the genes *AP3D1*, *RPL13A* and *PAK1IP1* are the strongest candidates overall. Our data suggests only a pair of genes is necessary for effective normalisation: *AP3D1* and one other. We and others have reported *RPL13A* to be a strong candidate under other scenarios [26, 37, 58–60], while literature for *PAK1IP1* is more limited: *AP3D1* plus *RPL13A* thus seems the most practical pair for normalising gene expression in the developing mouse embryo.

## Acknowledgements

This work was funded by the Wellcome Trust (Grant number 101550, awarded to RJP) and the Gill Malone Memorial Award (to JCWH). This manuscript has been approved by the RVC research office and assigned the following number: CSS_02293

## Author contributions

JCWH: Conceptualization, Formal Analysis, Funding acquisition, Investigation, visualization, Writing -original draft; DJW: Investigation, Resources, Supervision, Writing -review & editing; RJP: Funding acquisition, Supervision, Writing -review & editing.

