## Supplementary data for "Identification of qPCR reference genes suitable for normalising gene expression in the developing mouse embryo"

Reference gene analysis packages: methods, strengths and weaknesses

GeNorm

The geNorm algorithm [1] is an iterative pairwise approach: the method determines, for each gene in the panel, the stability value M (the mean pairwise variation with all other candidates). The gene with the highest M, i.e. the gene with expression behaviour least similar to all other candidates, is then discarded, and the analysis is repeated, discarding the gene with the highest M each time until only two genes with the lowest pairwise variation remain: the ‘best pair’. Assuming candidate genes are taken from unrelated cellular categories (thus unlikely to be co-regulated) two genes that exhibit closely-matched variation between samples are most likely to be reflecting differences in sample cDNA content. This method identifies shared patterns of variation, rather than stable expression *per se*, thus works well with comparatively noisy datasets, but is sensitive to extreme sample outliers. Furthermore, as a pairwise approach, the algorithm cannot identify a single candidate: each gene of the best pair has equal score (though the remaining genes can be ranked by their M values to provide an overall assessment of the panel). This algorithm is now integrated into the commercial qBase+ software package (Biogazelle, Zwijnaarde, Belgium - [www.qbaseplus.com](http://www.qbaseplus.com/)), however an archive copy of the original (non-commercial) excel macro is available at <http://ulozto.net/xsFueHSA/genorm-v3-zip>.

DeltaCt

As with geNorm, the deltaCt method [2] compares candidate genes against one another. This approach discards iterative comparisons in favour of a more direct assessment of candidate gene variability. For a given cDNA sample, the Cq value of a gene represents both the expression of that gene and the amount of cDNA present: comparing the difference in Cq values between two genes however eliminates [cDNA] as a variable, thus this ‘deltaCt’ reflects only the relative abundance of the two genes in that sample. If both genes are stably expressed, this difference should remain consistent across all samples, regardless of cDNA content. The standard deviation of deltaCt values between two genes thus reflects the stability for that specific pairing, and the mean of the deltaCt standard deviations for all possible pairings of a given gene represents that gene’s overall stability score, allowing the genes to be ranked accordingly. No dedicated software package is required for this method: deltaCt standard deviations can be derived manually from raw Cq values using Microsoft Excel.

Bestkeeper

Bestkeeper [3] takes the geometric mean of per-sample expression data for every gene in the dataset. The ‘bestkeeper’ expression profile generated by this method thus represents the mean behaviour of all candidate genes combined, with the rationale being that (for a sufficiently broad panel of genes) the average behaviour of all genes between samples should accurately reflect the cDNA content of those samples. By comparing the per-sample expression of each gene individually to that of this ‘bestkeeper’ (Pearson correlation, r), the gene or genes that serve as the best individual proxy for this consensus profile can be determined. A potential pitfall of this method is that an otherwise inappropriate gene might closely match the consensus profile by chance (as we have shown previously [4]), thus this method is best used in concert with others. The original software package (a write-protected Excel file) is available at <https://www.gene-quantification.de/bestkeeper.html>, however this package allows for only 10 candidate genes and 100 samples. We have prepared a simpler spreadsheet that allows larger panels (20 genes) and sample sizes (up to 400 samples), which is available on request. Ours does not offer the comprehensive functionality of the BestKeeper software package, but does derive a bestkeeper and calculate corresponding correlation coefficients for each gene (as used here).

Normfinder

Unlike the methods above, the approach used by the Normfinder algorithm [5] is not pairwise, instead assessing individual genes by their overall stability (consistency of expression) across the dataset. As such, the method is less suited to noisy datasets where substantial overall variability is present, but is tolerant of single extreme outliers. A powerful facet unique to this method is grouped analysis: samples can be allocated to user-defined groups before assessment. A gene that appears highly stable across the whole ungrouped dataset might exhibit modest but consistent bias when assessed in a grouped context. Moreover, within a grouped context this method can suggest a high scoring pair of genes, which may not include the highest scoring individual gene: two genes that exhibit consistent group-specific variation, but of opposite sign, can be combined to provide a highly stable average. Normfinder is an Excel Add-In, and is freely available at <https://moma.dk/normfinder-software>.
