## Supplementary Figure S1 for "Identification of qPCR reference genes suitable for normalising gene expression in the developing mouse embryo"

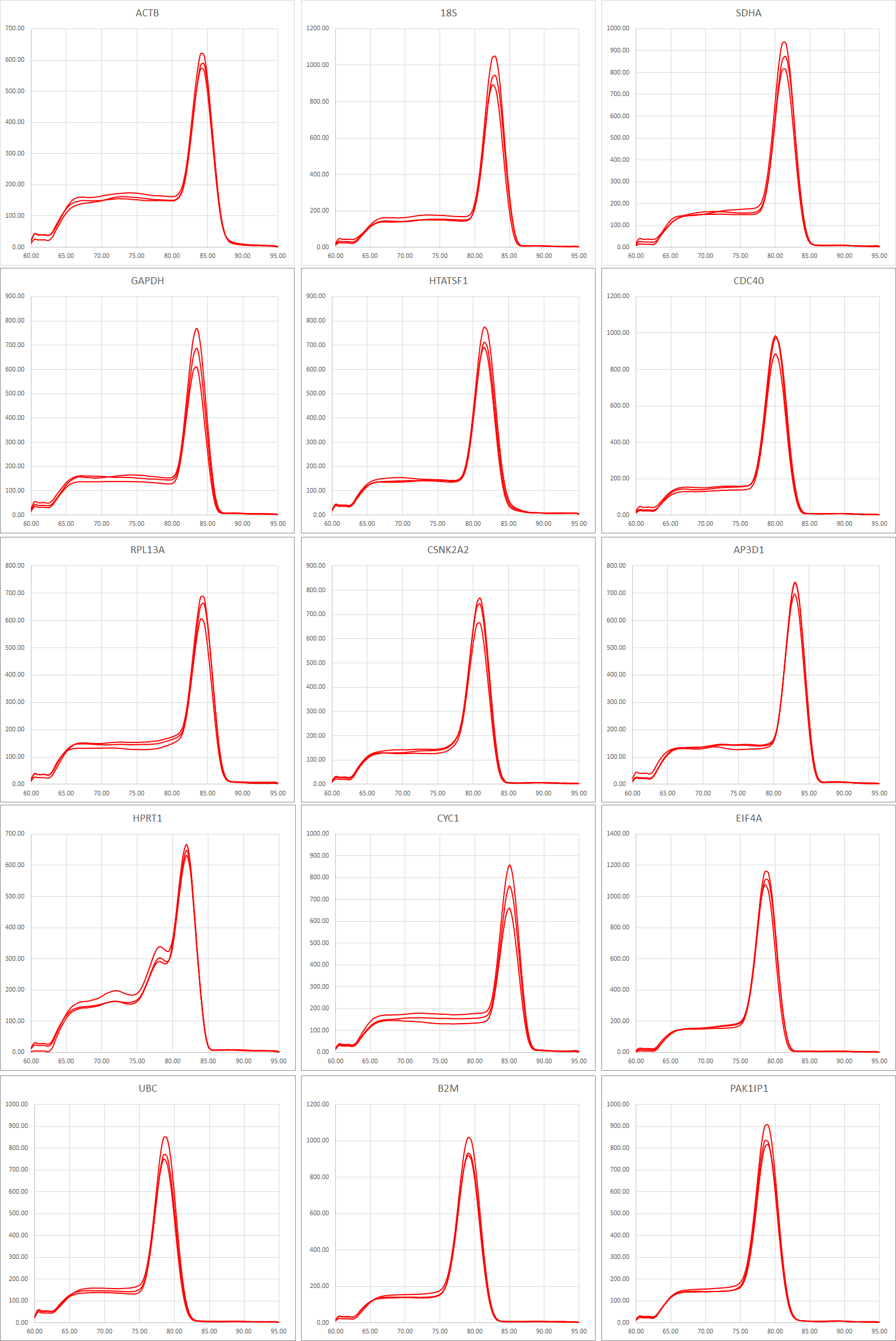

Supplementary Fig S1: amplicon melt curves

SYBR green melt curve derivatives for the amplicons produced by the fifteen primer pairs used in this study.
