## Supplementary Figure S2 for "Identification of qPCR reference genes suitable for normalising gene expression in the developing mouse embryo"

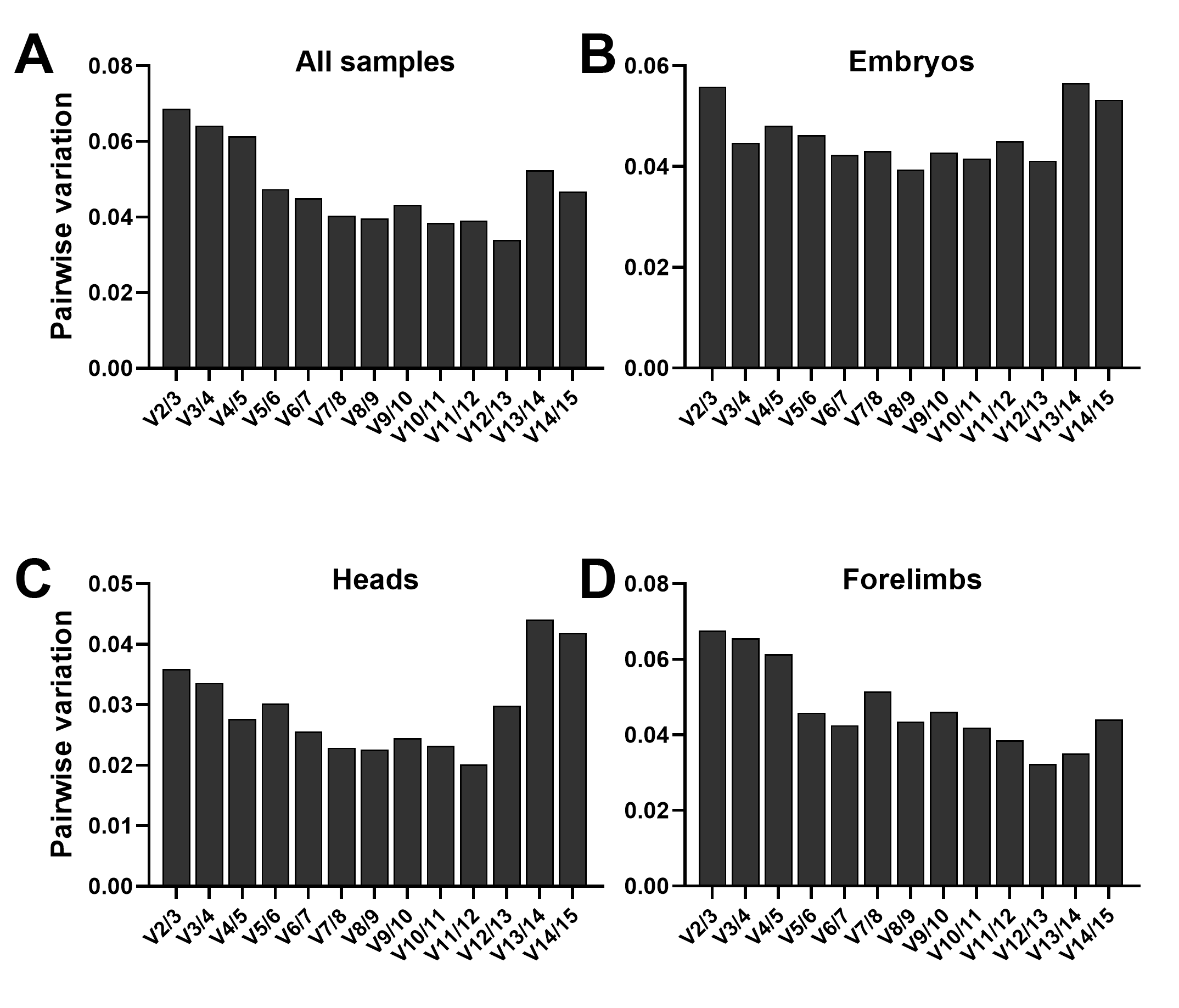

Supplementary Fig S2: additional geNorm analysis

GeNorm provides the change in pairwise variation elicited by increasing number of reference genes from two (best pair) to three or more as indicated. Values below 0.2 are considered sufficient. In all cases the best pair was sufficient, whether in our complete dataset (A), embryos alone (B), heads (C) or forelimbs (D).
