## Supplementary Figure S3 for "Identification of qPCR reference genes suitable for normalising gene expression in the developing mouse embryo"

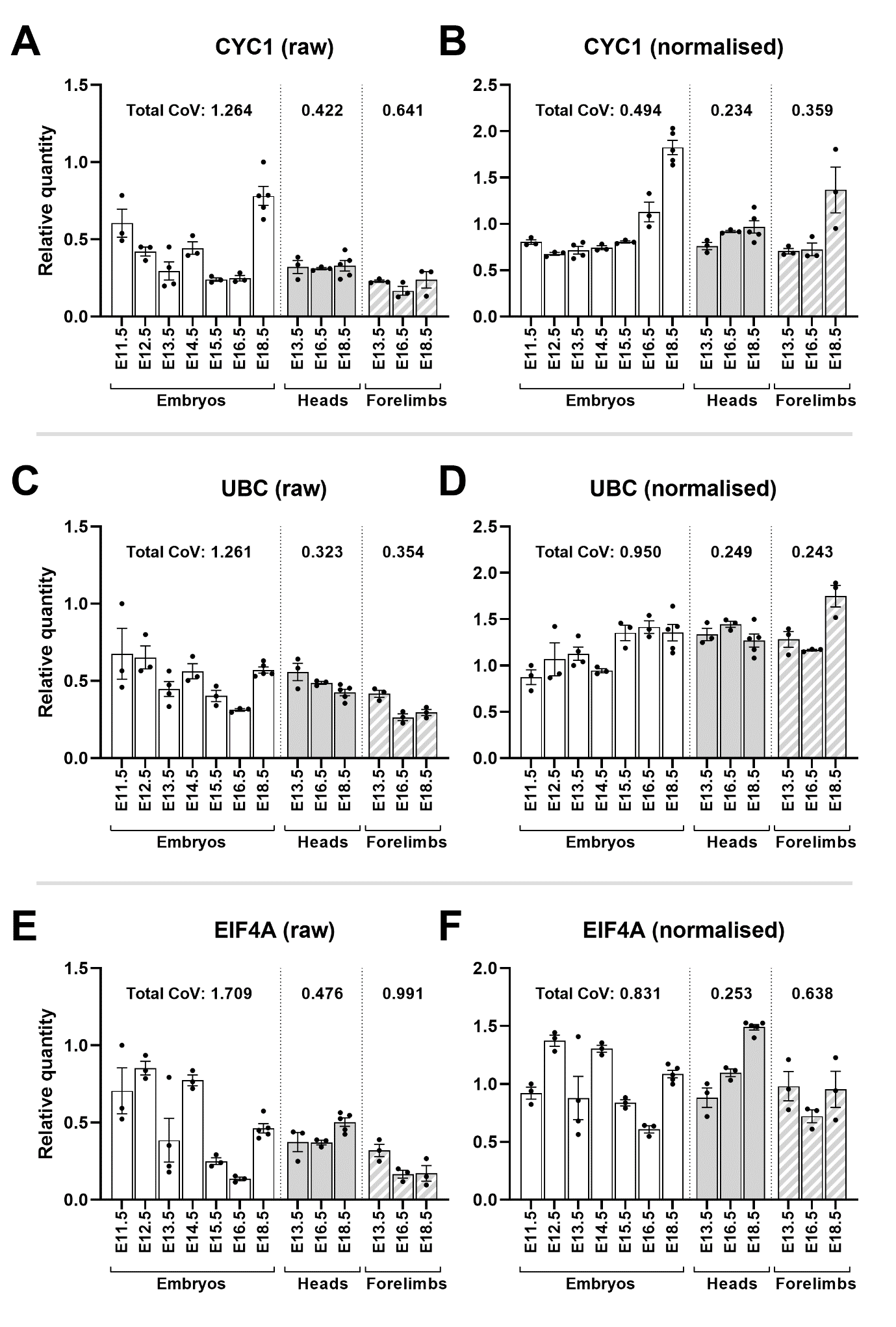

Supplementary Fig S3: additional validation: CYC1, UBC, EIF4A

Mean raw RQ values for CYC1 (A) and UBC (C) and EIF4A (E) are highly variable. Normalisation with AP3D1, RPL13A and PAK1IP1 (B, D and F) lowers total coefficient of variation (CoV) and reveals expression of CYC1 increases sharply at later developmental stages in whole embryos and forelimbs (but not in heads); UBC exhibits a similar but less dramatic upregulation, and EIF4A is variable in whole embryos, stable in forelimbs but is upregulated with gestational age in the head (as indicated). Total CoV values were obtained by summing the individual CoVs per time-point (see methods). Points represent individual RQ values (arbitrary units).
