## Supplementary Figure S4 for "Identification of qPCR reference genes suitable for normalising gene expression in the developing mouse embryo"

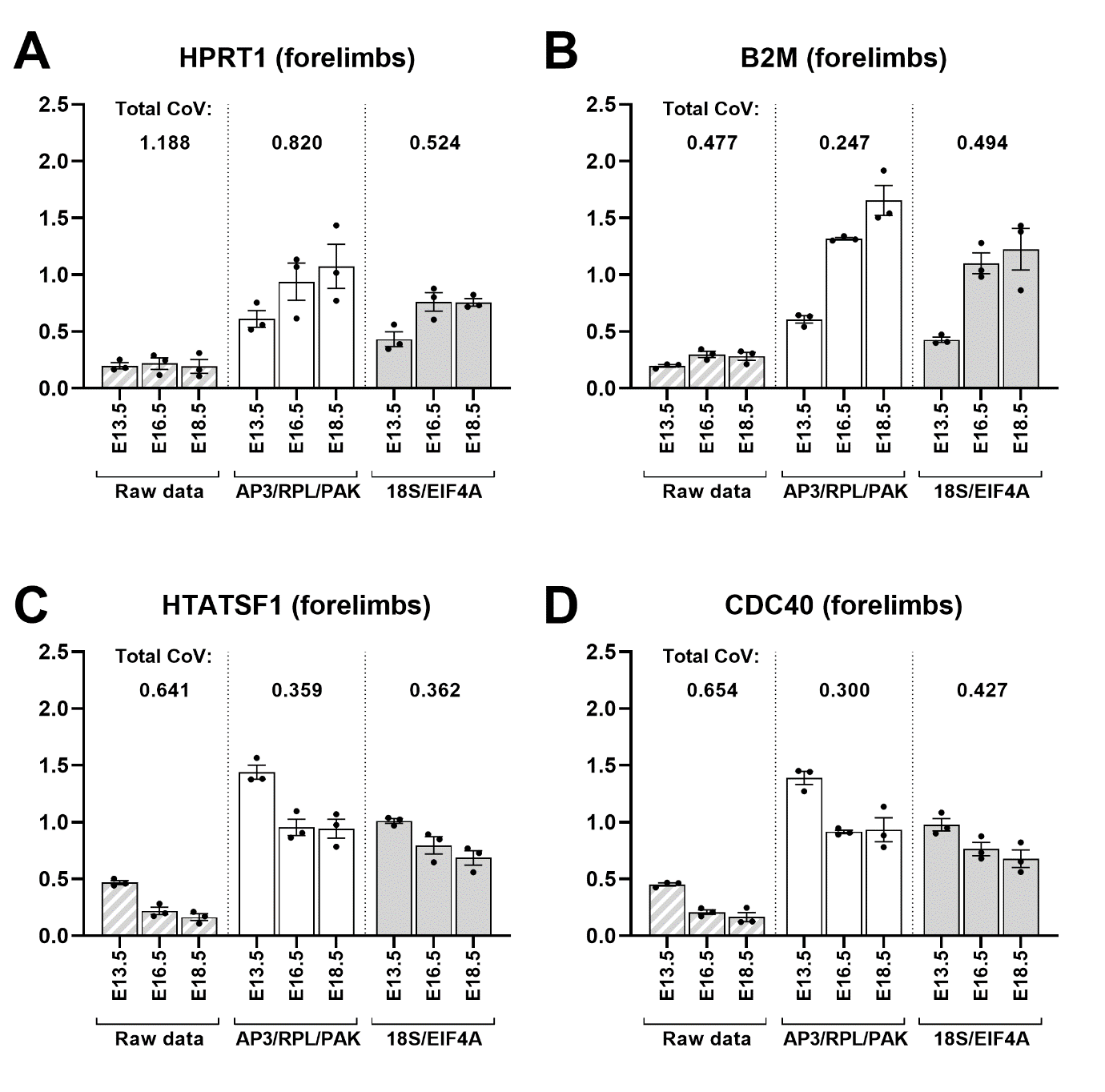

Supplementary Fig S4: AP3D1, PAK1IP1 and RPL13A are comparable to 18S and EIF4A in forelimbs

Mean RQ values for HPRT1 (A), B2M (B), HTATSF1 (C) or CDC40 (D), using raw data, or data normalised via the universally high-scoring AP3D1/RPL13A/PAK1IP1 or the forelimb specific high scoring 18S/EIF4A. Normalised data was comparable using either reference gene combination, and total coefficient of variation (CoV) was typically lower using the universal panel. Total CoV values were obtained by summing the individual CoVs per time-point (see methods). Points represent individual RQ values (arbitrary units).
