## Supplementary Figure S5 for "Identification of qPCR reference genes suitable for normalising gene expression in the developing mouse embryo"

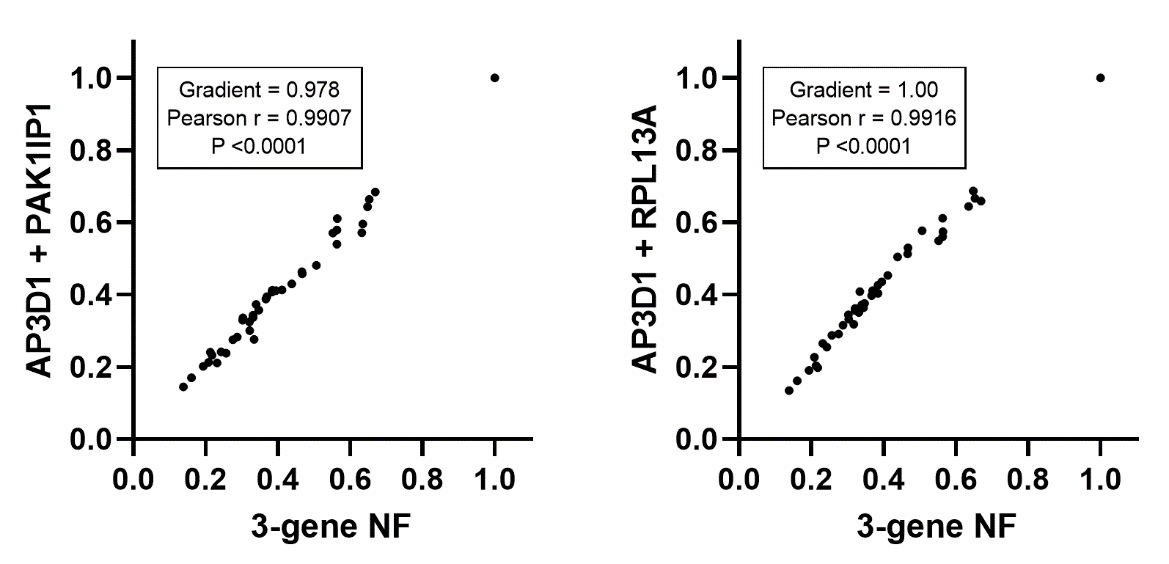

Supplementary Fig S5: Two reference genes are sufficient

Correlation plots of normalisation factors derived using AP3D1, RPL13A and PAK1IP1 (3-gene NF) with those derived using AP3D1 and only one other. In both cases use of two reference genes gave normalisation factors essentially indistinguishable from a 3-gene NF (gradients, Pearson correlations and P values as indicated).
