## Supplementary Table 1 for "Identification of qPCR reference genes suitable for normalising gene expression in the developing mouse embryo"

| Gestational age | Embryos  (total) | Used for this study | |
| --- | --- | --- | --- |
|  |  | Whole | Heads/forelimbs |
| 11.5 | 7 | 3 | - |
| 12.5 | 7 | 3 | - |
| 13.5 | 5 | 1 | - |
| 13.5 | 8 | 3 | 3 |
| 14.5 | 1 | 1 | - |
| 14.5 | 8 | 2 | - |
| 15.5 | 6 | 3 | - |
| 16.5 | 10 | 3 | 3 |
| 18.5 | 8 | 3 | 1 |
| 18.5 | 10 | 2 | 4 |

Supplementary Table S1: animal numbers and sample sizes

Embryos were collected at the indicated gestational ages (dpc): each row represents a single pregnant animal (10 in total), with total number of embryos collected (70), and numbers used for this study (35) listed accordingly.
